# Detecting inbreeding depression in structured populations

**DOI:** 10.1101/2023.08.14.552950

**Authors:** Eléonore Lavanchy, Bruce S. Weir, Jérôme Goudet

## Abstract

Measuring inbreeding as well as its consequences on fitness is central for many areas in biology including human genetics and the conservation of endangered species. However, there is no consensus on the most appropriate method, neither for quantification of inbreeding itself nor for the model to estimate its effect on specific traits. In this project, we simulated traits based on simulated genomes from a large pedigree and empirical whole-genome sequences of human data from populations with various sizes and structure (from the 1,000 Genomes project). We compare the ability of various inbreeding coefficients (*F*) to quantify the strength of inbreeding depression: allele sharing, two versions of the correlation of uniting gametes which differ in the weight they attribute to each locus and two identical-by-descent segments-based estimators. We also compare two models: the standard linear model and a linear mixed model including a genetic relatedness matrix (GRM) as random effect to account for the non-independence of observations. We find linear mixed models give better results in scenarios with population or family structure. Within the mixed models, we compare three different GRM matrices and show that in homogeneous populations, there is little difference among the different *F* and GRM for inbreeding depression quantification. However, as soon as strong population or family structure is present, the strength of inbreeding depression can be most efficiently estimated only if (i) the phenotypes are regressed on inbreeding coefficient based on a weighted version of the correlation of uniting gametes, which gives more weight to common alleles and (ii) with the GRM obtained from an allele sharing relatedness estimator.

## Introduction

Inbreeding is the result of mating between relatives and is often associated with reduced fitness, a phenomenon called inbreeding depression (ID) and which was observed in many different species such as humans [7, 6], other animals [26, 12, 21], and plants [34].

Many different methods have been developed for inbreeding quantification and there is no consensus on which one is the best [1, 5, 11, 25, 33, 35]. The classical approach was first proposed by Sewall Wright in 1922 and makes use of pedigrees (called hereafter *F*_PED_) [31]. With the advances in sequencing technologies, genomic-based inbreeding coefficients (hereafter called *F*_genomic_) have been developed. Among these, some coefficients rely on the comparison between observed and expected heterozygosity such as *F*_HOM_ [8, 27], the expected allele sharing between individuals such as *F*_AS_ [35] or on the correlation between uniting gametes such as *F*_UNI_ [32]. In addition to estimating the realized inbreeding coefficient and requiring no prior knowledge of the mating behavior of the population, these genomic estimates are simple and straightforward to compute and do not require whole-genome sequencing (WGS) data; a few thousands SNPs are usually sufficient for reliable inbreeding estimation in humans [11]. However they also have a disadvantage: they usually rely on allelic frequencies (except for *F*_AS_) and therefore if these frequencies have not been correctly estimated, this will affect the estimation of these coefficients. Another inbreeding coefficient was proposed by McQuillan *et al*. (2008): *F*_ROH_ uses runs of homozygosity (ROHs), long homozygous stretches as a proxy for IBD segments within individuals [22]. A model-based approach relying on hidden Markov models has also been developed for detecting IBD segments [19] by identifying homozygous-by-descent (HBD) segments. This model is the basis for many other model-based IBD segments detection methods such as `BCFTools` [24], `BEAGLE` [3] and `RZooRoH` [10]. The inbreeding coefficient estimated with these model-based approaches will be called *F*_HBD_ from now on. One advantage of these methods is that they do not depend on allelic frequencies which can be very valuable when only a few individuals are available. However, it has been shown that these coefficients, and especially *F*_ROH_, are sensitive to SNP density and parameters used, and there is no consensus on what is the most suitable set of parameters at present [23, 18].

How to quantify ID, although central to conservation genetics for decades [16], is still debated. This debate includes two sub-questions: which statistical model should be employed ? And which *F* ? Regarding the model, the classical approach consisted of the use of linear regression of the phenotypes on the inbreeding coefficient. However, other models have been utilized, such as Generalized Linear models (GLMs) with various link functions. In 2019, Nietlisbach *et al*.. [25] compared different models and found that the common GLM models with logit link did not allow for accurate inbreeding depression strength estimation. They propose using GLM with logarithm link functions. Ultimately, the type of model is largely dependent on the distribution of the trait.

Regarding the choice of which *F* is more accurate for quantifying ID, many studies have demonstrated that *F*_genomic_ yields better results than *F*_PED_ [17, 2, 13]. However, some studies found *F*_UNI_ to be more accurate than *F*_ROH_ [33], while others found that *F*_ROH_ provided the best estimates of ID [17, 13, 25]. In 2020, Caballero *et al*. [5] used simulations and included several populations with different histories: they found that the optimal *F* actually depends on how large the population is. *F*_ROH_ did a better job at quantifying ID in populations with small effective size while *F*_UNI_ was better at predicting ID estimates in populations with large effective sizes. This result was later confirmed by Alemu *et al*. [1] used SNP-array empirical cattle data for several groups of allelic frequencies and concluded that *F*_UNI_ and *F*_GRM_ (*F*_*I*_ and *F*_*III*_ respectively in [32]) are better at quantifying homozygosity at rare alleles while *F*_ROH_ and *F*_HOM_ are better for alleles at intermediate frequencies and correlate better with whole-genome homozygosity. Indeed, recessive deleterious alleles, which are thought to be responsible for inbreeding depression, should segregate at low frequencies in large populations as a result of negative selection. On the contrary, in small populations, drift can increase the frequency of deleterious recessive alleles to intermediate frequencies, making *F*_ROH_ and *F*_HOM_ more suitable for detecting ID. Indeed, in the simulations conducted by Yengo *et al*. [33], rare alleles always caused negative effects on fitness (referred to as DEMA, for Directional Effect of Minor Alleles). The authors showed that *F*_HOM_ (and thus *F*_AS_ since they have similar properties) is sensitive to DEMA while *F*_UNI_ and *F*_ROH_ are not. They also showed via simulations that all estimates of ID are somewhat sensitive to population structure, *F*_UNI_ being the least affected. They recommend estimating ID using Linkage Disequilibrium (LD) score and Minor Allele Frequency (MAF) bins, and summing the ID estimates from these bins as an overall estimate of ID for the trait.

In this paper we simulated traits based on both simulated and empirical WGS human data from populations with varying sizes and structure. We show that some *F* are more sensitive to population structure and DEMA than others. We confirm only some of Yengo *et al*. [33] results. Importantly, we show that accounting for the non-independence of observations with a mixed model via an allele sharing based genomic relationship matrix (GRM) (rather than the standard `GCTA` GRM) and using a modified version of *F*_UNI_ which gives more weight to common alleles resolves most of the issues raised by Yengo *et al*. [33].

## Material and Methods

### Simulated pedigrees

We simulated a polygamous pedigree from a dioecious population with overlapping generations (hereafter called PEDIGREE) using custom `R` scripts. The population started from 500 founders (equal numbers of males and females), and followed a polygamous mating system: female fertilities per time interval were drawn from a Poisson distribution with parameter *λ* = 1, mortality rate per time interval was set to 0.5, and only 10% of the males were allowed to reproduce at each time step. Matings were recorded for 25 time steps, resulting in a pedigree of 11, 924 individuals (over 25 time steps).

In order to simulate the genotypes of the individuals, we proceeded in two steps. We used the `mspms` wrapper to the `msprime` software [15] to simulate the two haplotypes containing *L* = 650, 000 loci for each founder individual. The *L* loci were uniformly distributed along a constant recombination map 20*M* long. For each reproduction event, the number of cross-overs was first drawn from a Poisson distribution and then randomly positioned along the genome. The non-founder genotypes were then obtained by drawing two gametes: one from each parent. For each gamete, the allele at the first locus is selected at random between the two alleles of the parent. The alleles at the next loci along the chromosome are copied from the chromosome with the chosen allele at the first locus until a recombination event occurs, at which point the alleles are copied from the other chromosome until the next crossing-over or the end of the chromosome.

In order to investigate the effect of using more realistic smaller sample sizes, we subsampled 2,500 individuals from the PEDIGREE population. We performed two types of sub-sampling: i) a random sub-sampling where individuals were subsampled completely randomly, ii) a stratified sub-sampling where we sought to retain the widest range of inbreeding coefficients in the sub-sampled population. Consequently, for this stratified sub-sampling individuals with 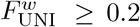 were always included and individuals with 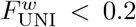 were randomly selected until the population reached the desired size. 100 replicates were performed for each sub-sampling.

### 1000 Genomes

In order to extend our conclusions to even smaller sample sizes and populations with stronger structure (which are common in wild and/or endangered species), we used empirical data from phase 3 from the 1,000 Genomes project [28]. We considered i) a small sample from a homogeneous population with small effective size represented by 504 individuals from the super-population with East-Asian ancestry (EAS), ii) a small sample from a population with some admixture and larger effective population sizes represented by 661 individuals from the super-population with African ancestry and admixed individuals (AFR) and finally iii) a larger sample from a population with larger effective size and with genetic structure (global *F*_*ST*_ = 0.083) comprising all the 2,504 individuals (hereafter called WORLD) and represented by five super-populations: individuals with East-Asian ancestry (EAS), African ancestry (AFR), European ancestry (EUR), admixed American ancestry (AMR) and finally South-Asian ancestry (SAS). A more detailed description of the samples can be found at the [1,000 Genomes Project website](https://www.internationalgenome.org/).

### Simulated traits

We simulated traits based on equation 1 following [33]: we consider a trait *y* whose phenotype is partly determined by the genotypes at *Lc* causal loci with *h*^2^ = 0.8. We assume these loci to be bi-allelic, with one allele encoding for an increase in the trait value (the plus allele) and the other encoding for a decrease in trait value (the minus allele). Dominance was also considered since inbreeding depression (ID) occurs only if there is directional dominance: when heterozygotes at loci encoding for the trait are closer on average to the homozygote for the plus allele [20]. If gene effects are purely additive or if dominance is not directional, there is no ID. Finally, we assume no epistasis between loci, and no genotype-environment interaction.

For individual *j, y*_*j*_ is the individual trait value (its phenotype), calculated as the sum of allelic and genotypic effects over causal loci, an environmental effect and *μ*, the average trait value among all individuals. At locus *l, x*_*jl*_ is the minor allele count (MAC) ∈ *{*0, 1, 2*}* of individual *j. a*_*l*_ represents the additive effect size of the alternate allele at locus *l. d*_*l*_ is the dominance effect size, the deviation of the heterozygous genotype from the mean of the two homozygotes. Finally, *ϵ*_*j*_ is the environmental contribution to the phenotype of individual *j*, drawn from a normal distribution.

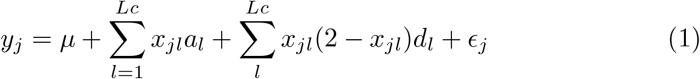

The strength of inbreeding depression *b* was set to −3 in all simulations, as in Yengo *et al*. [33]. We chose a value which was close to zero because if the the effect of inbreeding is too strong, it will always be detected. In addition, this value is in the range of observed inbreeding depression published estimates (for instance, table 10.4 from [20]).

We used equation 1 to simulate traits with varying architectures. To avoid causal markers with extremely low frequencies, we first excluded loci with *MAF* ≤ 0.01 for both the EAS and AFR populations and loci with *MAF* ≤ 0.001 for both the PEDIGREE and WORLD populations. We then simulated traits using 1,000 randomly chosen SNPs (after MAF filtering). We drew both the raw additive effect sizes of the alternate allele and the raw dominance effect sizes from a uniform [0, 1] distribution (other distributions were explored with almost no effect on the results (results not shown)). As we expect alleles causing ID to be counter selected and thus removed or maintained at a low frequency (proportionally to their detrimental effect), the raw effect sizes were scaled inversely to MAF *a*_*j*_ = *raw*_*aj*_*/p*_*j*_ to mimic negative selection. We also scaled the dominance effects inversely to the locus expected heterozygosity *d*_*j*_ = *raw*_*dj*_*/*(2*p*_*j*_(1 − *pj*)). In addition, we attributed the same sign to the effect sizes of all minor alleles in order to include what Yengo *et al*. [33] called Directional Effect of Minor Alleles (DEMA) [33]. However, in order to investigate the effect of the parameters mentioned above, we also simulated traits where the additive and dominance effect sizes were left unchanged *a*_*j*_ = *raw*_*aj*_ and *d*_*j*_ = *raw*_*dj*_ and without DEMA. A summary of all the simulated scenarios can be found in table S1. In addition, graphical representation of the additive effect sizes and dominance coefficients distribution under these different scenarios can be found in Figure S1.

### Individual inbreeding coefficients

We estimated individual inbreeding coefficients using several methods whose properties were recently described in detail in Zhang *et al*. [35]. Regarding the figures and tables presented in the main text, we do not filter on MAF for any of the *F* s estimates. We use one allele-sharing-based estimator of inbreeding, hereafter called *F*_AS_ and described in [30, 35]:

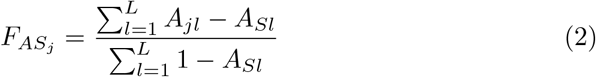

where *A*_*jl*_ indicates the identity of the two alleles an individual *j* carries at locus *l*: one for homozygous and 0 for heterozygous and *A*_*Sl*_ is the average allele sharing proportion at locus *l* for pairs of individuals *j, k, j* ≠ *k*.

Then, we compare two versions of *F*_UNI_ (initially described in [32]) and which measure the correlation between uniting gametes. The first version (hereafter called 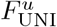) is the original *F*_UNI_ [32] measured as the average of ratios over SNPs (which attributes equal weight to all loci):

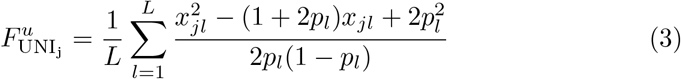

Similarly to equation 1, *x*_*jl*_ is the MAC of individual *j* at locus *l* ∈ *{*0, 1, 2*}* and *p*_*l*_ is the derived allele frequency at locus *l*.

The second version (hereafter called 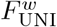) is a modified version of *F*_UNI_ which measures the ratio of averages and thus gives more weight to loci with larger expected heterozygosity (i.e. with MAF close to 0.5). We are not aware of other investigations using the ratio of averages estimator 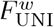 in the context of ID estimation.

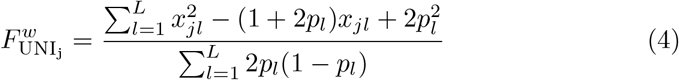

We also used four Identical-by-descent (IBD) segments based *F*. We called runs of homozygosity (ROHs) with `PLINK` [27] and default parameters. We also called Homozygous-by-descent (HBD) segments with `BCFTools` [24]. For both methods, we selected ROHs or HBD segments based on their size: either larger than 100Kb: 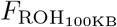 and 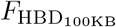 or larger than 1Mb: 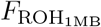 and 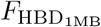. For both methods the inbreeding coefficients were simply estimated as the fraction of genome falling within ROHs or HBD segments.

Finally, in the PEDIGREE population, we used the pedigree-based inbreeding coefficient: *F*_PED_ [31].

All inbreeding coefficients were estimated separately for each population of the 1,000 Genomes Project (EAS, AFR, WORLD) and with population specific SNPs and allelic frequencies (i.e. we removed monomorphic SNPs and estimated allelic frequencies in all the three populations). Consequently the same individual might have different *F*_genomic_ in the EAS and the WORLD population. This influenced only the IBD segments-based inbreeding coefficients (*F*_ROH_ and *F*_HBD_) trivially but greatly influenced *F*_AS_ (though the rank of inbreeding among individuals was conserved) and both *F*_UNI_ (for which the rank of inbreeding among individuals was not conserved) since their formulae rely on allelic frequencies estimations. Comparison among the different inbreeding coefficients per population can be found in supplementary material (Figures S2 - S5). More details can be found in [35],

### Estimation of Inbreeding Depression: *b*

We estimated the strength of ID (hereafter defined as *b*) using two different models. In the first model, *b* was estimated as the slope of regression of phenotypes on the different inbreeding coefficients with a classical linear model (LM):

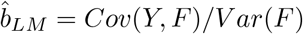

where *Y* is the vector of trait values and *F* is the vector of individual inbreeding coefficients estimates.

In the second model, we estimate *b* as the fixed effect coefficient associated with the inbreeding coefficient in the following linear mixed model (LMM):

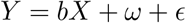

where *Y* is the vector of trait values, *X* is a matrix with two columns, the first containing ones and the second the individual inbreeding coefficients, *ω* is the random component of the mixed model with *ω* ∼ *N* (0, *τ K*), *K* being the genomic relationship matrix (GRM) and *τ* the additive variance component. Finally, *ϵ* is the individual residual variance and is defined as *ϵ* ∼ *σ*^2^*I*_*n*_. From this, *b* is estimated as follows:

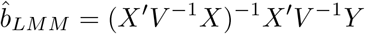

with *V* = *τ K* + *σ*^2^*I*_*n*_ [9]. We compare three different GRMs we estimated using all loci (no MAF filtering). The first mixed model included a GRM derived from allele sharing [11], hereafter called LMM_AS_. We used the `R Hierfstat` package to estimate *K* and the `R gaston` package to estimate *V* and *b*. We could not use `GCTA` software to run the mixed model for this GRM because its leading eigenvalue is negative which the Choleski decomposition algorithm used for matrix inversion in `GCTA` cannot handle (it requires a positive definite matrix), while the Schur decomposition algorithm used in `gaston` can. We note that the standard GRM is not positive definite (one eigen value is 0), but the matrix to invert in the mixed model is not the GRM itself but *V* = *τ K* + *σ*^2^*I*_*n*_ which becomes positive definite and can be inverted if the heritability is smaller than one.

The second mixed model used the `GCTA` weighted GRM matrix [11, 29]. Similarly to 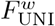, this matrix uses the ratio of averages. For this model, we used `GCTA` and the `R SNPrelate` package to estimate *V*. We then used the `R` `gaston` package for estimating *b* with the LMM.

Finally, the third mixed model used the `GCTA` unweighted GRM matrix [32] which (similarly to 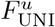) utilizes the average of ratios and thus gives equal weight to all loci. For this model, we used `GCTA` to estimate *V*. We then estimated *b* with the LMM implemented in the `R gaston` package.

Note that the Average Information-Restricted Maximum Likelihood (AIREML) fitting method we used in the LMM is an iterative procedure, and should result in unbiased estimates. In some cases, the model did not converge, and gave highly biased *b*. For each scenario, regression model and population, the number of replicates which did not converge can be found in tables S6-S8.

## Results

All the figures presented in the main text picture the scenario where alleles additive effect sizes and dominance coefficients are proportional to MAF and where there is a directional effect of minor alleles (DEMA) (i.e. the ADD & DOM & DEMA scenario from table S1) (see Figure S1). The results for the other scenarios are shown and discussed in supplementary material (Figures S8-S15, tables S2-S5).

### Simulated pedigrees

Figure 1 presents the inbreeding depression (ID) strength estimates (*b*, see the methods section) for the different inbreeding coefficients (*F*), with two regression models in the PEDIGREE populations. The first column shows *b* estimated with the simple LM and the second column shows *b* estimated with LMM including the allele sharing GRM as random factor (LMM_AS_). The first row shows results for the complete PEDIGREE population (n = 11,924). The second row shows results for a reduced sample size of the PEDIGREE population (n = 2,500, meant to match the size of the 1KG WORLD population) where sub-sampled individuals were chosen completely randomly. The third row also shows results for a reduced sample size of the PEDIGREE population (n = 2,500) but these individuals were selected to represent the entire spectrum of inbreeding values. The violin plots show *b* estimates distributions among the simulation replicates (100 replicates for the complete population, 10,000 replicates for both sub-sampled populations). The solid dark grey line is the true strength of ID (*b* = -3). The dashed red line represents the absence of ID (*b* = 0), indicating that ID was not detected in any replicate above this line. Root mean square error (RMSE) values associated with both models and populations are shown in table 1. Strikingly, in the PEDIGREE population, no *F* resulted in a accurate estimation of *b* with the simple LM, whatever the sample size (Figure 1, panels A, C and E; Table 1). The inclusion of a GRM matrix as a random factor allowed for the correction of non-independence of observations and greatly improved *b* estimation (Figure 1, panels B, D, and F; table 1). In the complete PEDIGREE population, we see little difference between the three GRMs we tested (1, panel B vs Figure S8, panels A and B; table 1): all *F* yielded efficient (we use efficient to describe an estimate with low RMSE, thus which is unbiased and has low variance) estimates of *b* when used inside a LMM, except for 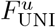 that slightly overestimates the strength of ID while *F*_PED_ slightly underestimates it. This suggests that large sample sizes (here 11,924 individuals) combined with a mixed model allow efficient ID estimation regardless of the *F* used. The three mixed models, however, perform less efficiently when the sample size is reduced, as we demonstrate with both subsampled PEDIGREE populations (n = 2,500): many replicates produced estimates above zero for *b* (Figure 1, panels D and F; Figure S8, panels C to F; table 1). RMSE were particularly large for *F*_PED_, 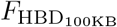 and 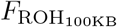 with the mixed model using the unweighted `GCTA` GRM matrix 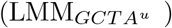 (Figure S8, panel D; table 1). Additionally, increasing the variance of sub-sampled individuals’ *F* (i.e. ranged sub-sampling) led to better estimates of *b* with reduced variance among replicates compared to random subsampling (Figure 1, panels D vs F: Figure S8, panels C vs E and D vs F, table 1).

**Table 1:** RMSE on *b* estimate in the PEDIGREE population. These values are for the complete ADD & DOM & DEMA scenario. See tables S2-S5 for the other scenarios

| Model | Population | $F_{\text{PED}}$ | $F_{\text{AS}}$ | $F_{\text{UNI}}^u$ | $F_{\text{UNI}}^w$ | $F_{\text{HBD}_{100\text{KB}}}$ | $F_{\text{ROH}_{100\text{KB}}}$ | $F_{\text{HBD}_{1\text{MB}}}$ | $F_{\text{ROH}_{1\text{MB}}}$ |
| --- | --- | --- | --- | --- | --- | --- | --- | --- | --- |
| LM | PEDIGREE (complete) | 34.82 | 22.71 | 10.17 | 4.17 | 19.93 | 22.22 | 17.4 | 17.44 |
| LMM: AS | PEDIGREE (complete) | 1.62 | 1.27 | 1.89 | 0.87 | 1.07 | 1.12 | 1.11 | 1.11 |
| LMM: GCTA WE | PEDIGREE (complete) | 1.62 | 1.27 | 1.89 | 0.87 | 1.07 | 1.12 | 1.11 | 1.11 |
| LMM: GCTA UN | PEDIGREE (complete) | 1.58 | 1.28 | 1.85 | 0.88 | 1.08 | 1.12 | 1.08 | 1.08 |
| LM | PEDIGREE (random sub) | 33.84 | 22.20 | 10.41 | 4.47 | 19.53 | 21.72 | 17.24 | 17.28 |
| LMM: AS | PEDIGREE (random sub) | 4.01 | 2.97 | 3.82 | 1.83 | 2.57 | 2.73 | 2.56 | 2.57 |
| LMM: GCTA WE | PEDIGREE (random sub) | 4.01 | 2.97 | 3.82 | 1.83 | 2.57 | 2.73 | 2.56 | 2.57 |
| LMM: GCTA UN | PEDIGREE (random sub) | > 1,000 | 2.75 | 3.44 | 1.78 | > 1,000 | > 1,000 | > 1,000 | > 1,000 |
| LM | PEDIGREE (ranged sub) | 15.22 | 11.04 | 3.46 | 1.61 | 9.58 | 10.52 | 8.13 | 8.15 |
| LMM: AS | PEDIGREE (ranged sub) | 2.09 | 1.82 | 2.13 | 1.26 | 1.61 | 1.67 | 1.58 | 1.58 |
| LMM: GCTA WE | PEDIGREE (ranged sub) | 2.09 | 1.82 | 2.13 | 1.26 | 1.61 | 1.67 | 1.58 | 1.58 |
| LMM: GCTA UN | PEDIGREE (ranged sub) | > 1,000 | 1.69 | 2.05 | 1.24 | > 1,000 | > 1,000 | 1.53 | 1.54 |

**Figure 1:**
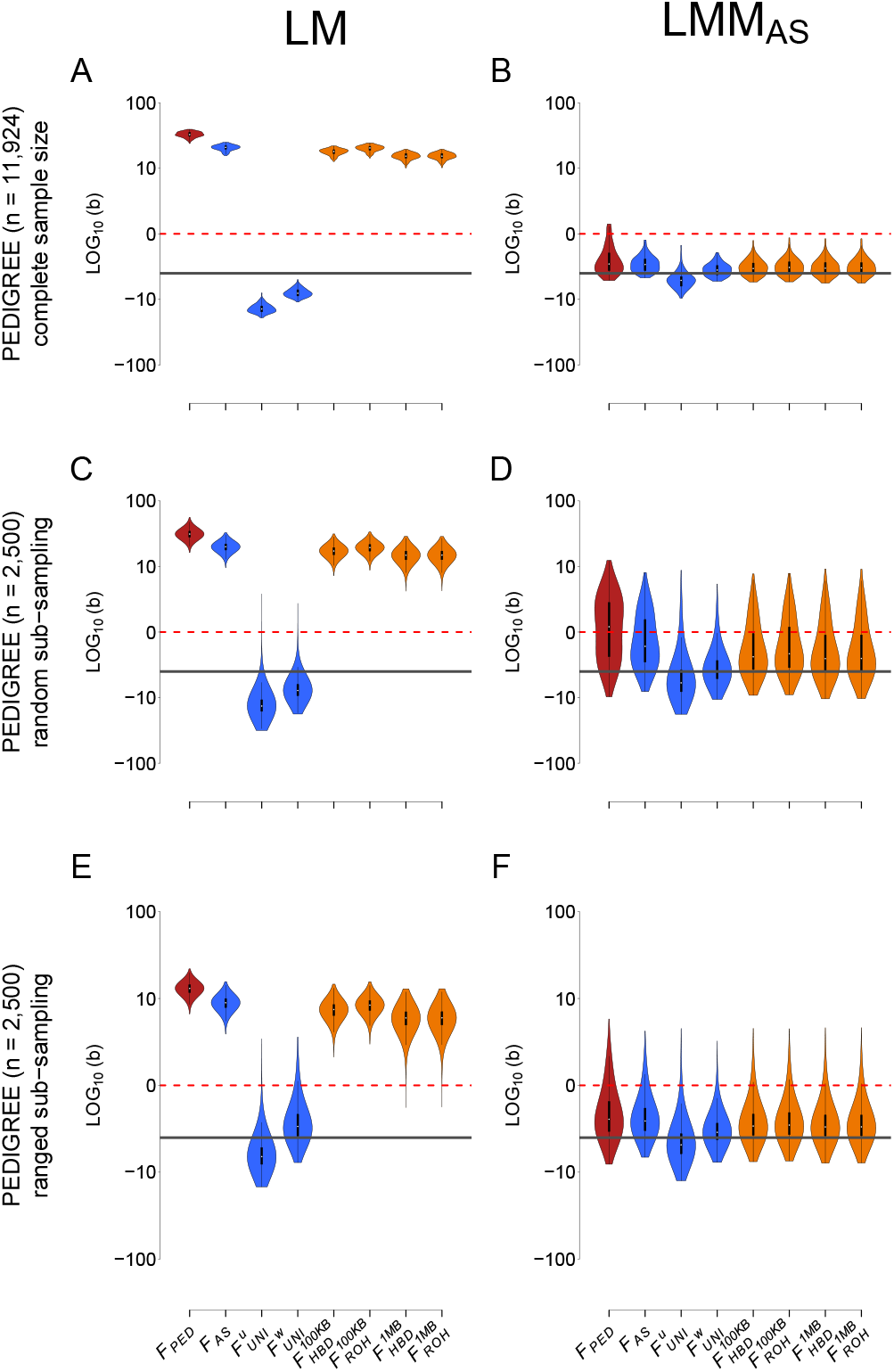
**Comparison of the estimation of inbreeding depression strength (***b*) among different *F* estimates and two models in the PEDIGREE population. Each column represents a regression model. The first column depicts the simple linear regression (LM) (panels **A, C** and **E**) and the second column depicts the linear mixed model with the allele sharing relatedness matrix as a random component (LMM_AS_) (panels **B, D** and **F**). The first row represents the complete simulated population (11,924 individuals, panels **A** and **B**). The second row shows the random subsampling (2,500 individuals, panels **C** and **D**). The third row shows the ranged subsampling (2,500 individuals, panels **E** and **F**). Inbreeding estimates presented in this graph are *F*_PED_, *F*_AS_, 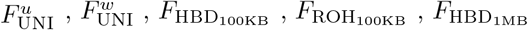 and finally 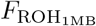. For panels **A** and **B**, violin plots show the distribution of the inbreeding depression strength estimates (*b*) among the simulated 100 replicates. For panels **C** to **F**, violin plots represent the distribution of the inbreeding depression strength estimates (*b*) for the 10,000 simulated and sub-sampling replicates (100 sub-sampling replicates for each of the 100 simulation replicates). The solid dark grey line is the true strength of ID (*b* = -3). The dashed red line represents the absence of ID (*b* = 0), meaning that we failed to detect ID in any replicate above this line. Note that all panels are in *log*10 scale.

### 1,000 Genomes Project

Figure 2 illustrates the estimates of ID strength (*b*) for the different inbreeding coefficients (*F*), when using either a LM or a LMM for two subsets of the 1,000 Genomes Project: EAS and AFR, as well as for the entire world population. It has the same structure as Figure 1. Root mean square error (RMSE) values associated with both models and populations can be found in table 2. Interestingly, we see little difference between LM and LMM and the different GRMs when there is no structure among the samples even with small sample sizes (**EAS**: Figure 2, panel A and B vs Figure S6, panels A and B; table 2; **AFR**: Figure 2, panel C and D vs Figure S6, panels C and D; table 2). Similarly to what was observed for the PEDIGREE population, when some structure exists (population structure in the WORLD population compared to family structure in the PEDIGREE population), the simple LM fails to accurately estimate the strength of ID, regardless of the *F* (Figure 2, panel E; table 2). In contrast to the pedigree population showing no difference between the three GRMs (Figure 1 and Figure S6), the most efficient estimates of *b* are obtained only with the LMM_AS_ model and with 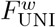 in the highly structured WORLD population (Figure 2, panel F vs Figure S7 panels E and F; table 2). In fact, the models including the *GCTA*^*w*^ and *GCTA*^*u*^ matrices cannot efficiently estimate *b* with any of the inbreeding coefficients: even though 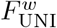 is unbiased, the variance is very large (panel F; Figure S7, table 2). In addition, several replicates did not converge when both *GCTA*^*w*^ and *GCTA*^*u*^ models were used which was never the case with the *GRM*_AS_. Numbers of such replicates are indicated in the Figures’ legend and in supplementary tables S6-S8.

**Table 2:** RMSE on *b* estimate in the three 1,000 Genomes Project populations: EAS, AFR and WORLD. These values are for the complete ADD & DOM & DEMA scenario. See tables S2-S5 for other scenarios

| Model | Population | $F_{AS}$ | $F_{UNI}^u$ | $F_{UNI}^w$ | $F_{HBD_{100KB}}$ | $F_{ROH_{100KB}}$ | $F_{HBD_{1MB}}$ | $F_{ROH_{1MB}}$ |
| --- | --- | --- | --- | --- | --- | --- | --- | --- |
| LM | EAS | 5.55 | 4.9 | 4.86 | 7.14 | 7.93 | 6.19 | 10.58 |
| LMM: AS | EAS | 5.67 | 4.68 | 4.64 | 7.41 | 8.22 | 6.12 | 10.39 |
| LMM: GCTA WE | EAS | 5.67 | 4.68 | 4.64 | 7.28 | 8.06 | 6.11 | 10.39 |
| LMM: GCTA UN | EAS | 5.48 | 4.74 | 4.71 | 7.1 | 7.87 | 6.18 | 10.57 |
| LM | AFR | 5.93 | 4.81 | 4.81 | 6.03 | 7.21 | 7.21 | 13.12 |
| LMM: AS | AFR | 5.15 | 4.07 | 4.07 | 5.46 | 6.2 | 7.15 | 13.1 |
| LMM: GCTA WE | AFR | 5.15 | 4.07 | 4.07 | > 1,000 | > 1,000 | 7.16 | 13.1 |
| LMM: GCTA UN | AFR | 5.78 | 4.42 | 4.42 | 5.92 | 6.93 | 7.2 | 13.11 |
| LM | WORLD | 32.91 | 142.95 | 62.21 | 67.42 | 59.15 | 107.67 | 169.73 |
| LMM: AS | WORLD | 8.63 | 8.34 | 4.17 | 9.15 | 10.97 | 8.78 | 14.6 |
| LMM: GCTA WE | WORLD | 9.84 | > 1,000 | > 1,000 | 11.19 | 13.92 | > 1,000 | > 1,000 |
| LMM: GCTA UN | WORLD | 18.18 | > 1,000 | > 1,000 | 27.52 | 26.91 | > 1,000 | > 1,000 |

**Figure 2:**
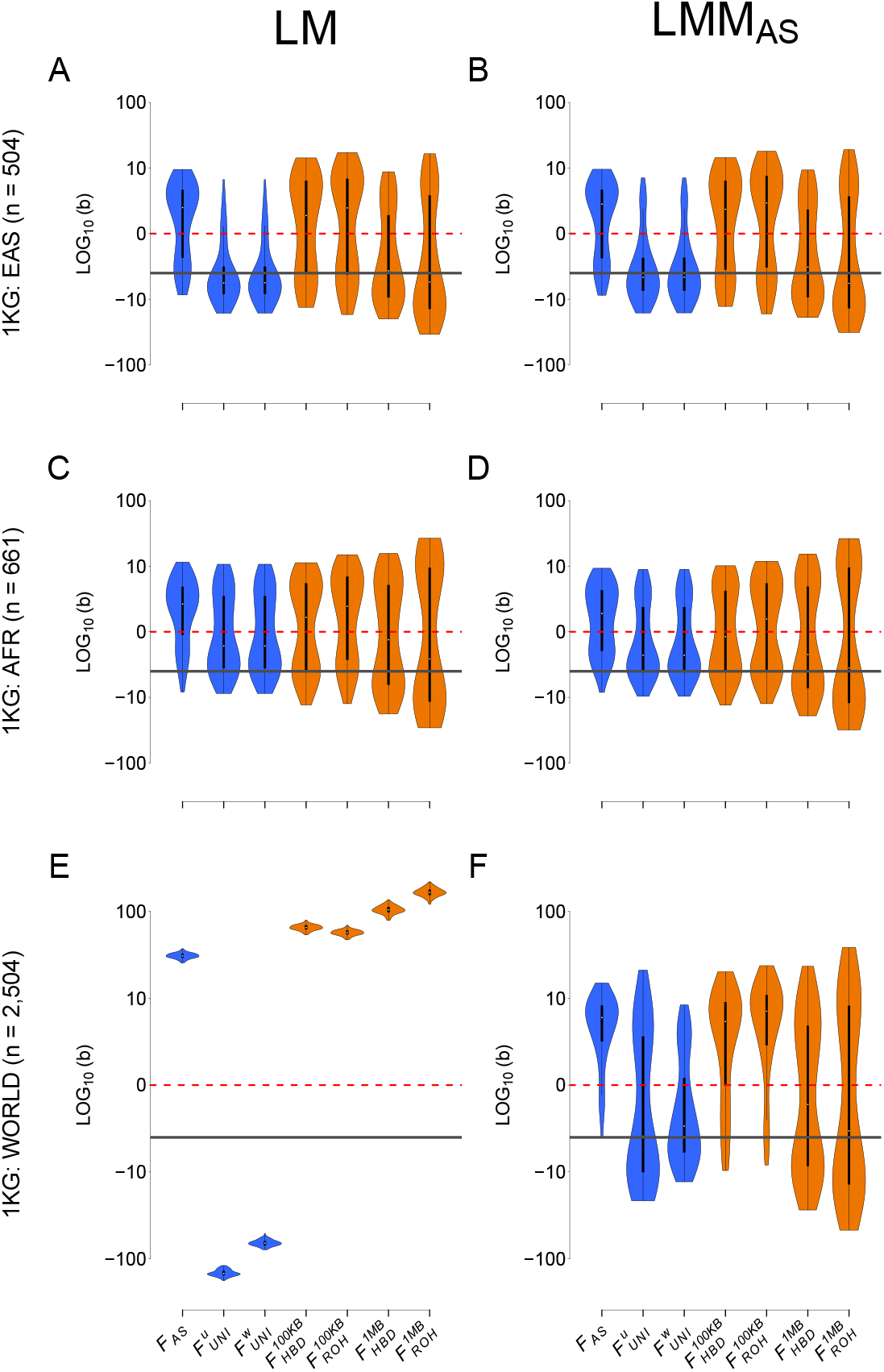
**Comparison of the estimation of inbreeding depression strength (***b*) among different *F* estimates and models in four different populations. Each column represents a regression model. The first column depicts the simple linear regression (LM) (panels **A, C** and **E**) and the second column depicts the linear mixed model with the allele sharing relatedness matrix as a random component (LMM_AS_) (panels **B, D** and **F**). The three rows correspond to the three populations from the 1,000 Genomes project: EAS on panels **A** and **B**, AFR on panels **C** and **D** and WORLD on panels **E** and **F**. Inbreeding estimates presented are *F*_AS_, 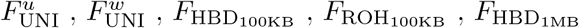 and finally 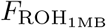. Violin plots represent the distribution of the inbreeding depression strength estimates (*b*) among the 100 simulation replicates. The solid dark grey line is the true strength of ID (*b* = -3). The dashed red line represents the absence of ID (*b* = 0), meaning that we failed to detect ID in any replicate above this line. Note that all panels are in *log*_10_ scale.

### Comparing inbreeding coefficients

With both the LM and LMM_AS_ models in the three populations from the 1,000 Genomes Project (EAS, AFR and WORLD, panels A - F) and for the LM in the PEDIGREE population, *F*_AS_ is consistently underestimating the strength of ID, particularly when there is strong structure (WORLD: Figure 2, panels E and F). It is because DEMA is included in the model and strongly influences the quantification of ID by *F*_AS_. In the absence of a DEMA, *F*_AS_ produces efficient estimates (Figures S10 - S13). In addition, *F*_AS_ is sensitive to the dominance effects being proportional to MAF but to a lesser extent and in the opposite direction (Figure S8 vs Figure S9). Concerning the other SNP-based *F*, 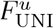 is constantly overestimating the strength of ID and is the most sensitive to population structure: its variance is much larger compared to 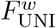 in the structured WORLD population and with all models (Figure 2, panel F; table 2). Interestingly, the variance of 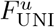 is affected only when allele effect sizes and/or dominance coefficients are proportional to MAF, but not by DEMA (Figures S8-S15). In contrast, 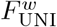 is the least sensitive to allele effect sizes or dominance coefficients proportional to MAF and DEMA (Figures S8 – S15), which makes it the most appropriate *F* for estimating ID (Figure 2, panel F; table 2). Since the difference between 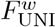 and 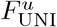 is the weight given to rare and common alleles, we conducted the same analyses (including the re-estimation of both *F* and GRMs estimation) on the WORLD population but excluding loci with MAF *>* 0.05 and showed that there is no difference between 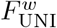 and 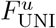 when rare alleles are removed (Figure S16). Concerning the *F* calculated from ROHs and HBD segments, there is not much difference between `PLINK` and `BCFTools` except for the variance among *b* estimates, which is slightly smaller with `BCFTools` compared to `PLINK` (Figure 2, panels A - F; table 2). In addition, focusing on recent inbreeding by including only large segments (here larger than 1MB) yielded better results in the WORLD population (Figure 2, panel F). Since `BCFTools` is a model-based HBD approach, there is no mandatory length requirement. In light of this, we also estimated *F*_HBD_ based on HBD segments without any size restrictions, and the results are similar to those obtained using 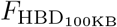 (Figure S17).

### Comparing genetic relatedness matrices

Since we identified 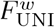 as the best inbreeding coefficient, Figure 3 contrasts the four different models for this coefficient in the four populations: each panel corresponds to one population. As mentioned above, there is almost no difference among the different GRM matrices in the extremely large complete PEDIGREE population (Figure 3, panel A; table 1) and between any of the models in the two homogeneous populations (EAS and AFR) (Figure 3, panels B and C; table 2). However, in the highly structured WORLD population, LMM_AS_ gives the most accurate result due to its smaller variance and RMSE (Figure 3, panel D; table 2).

**Figure 3:**
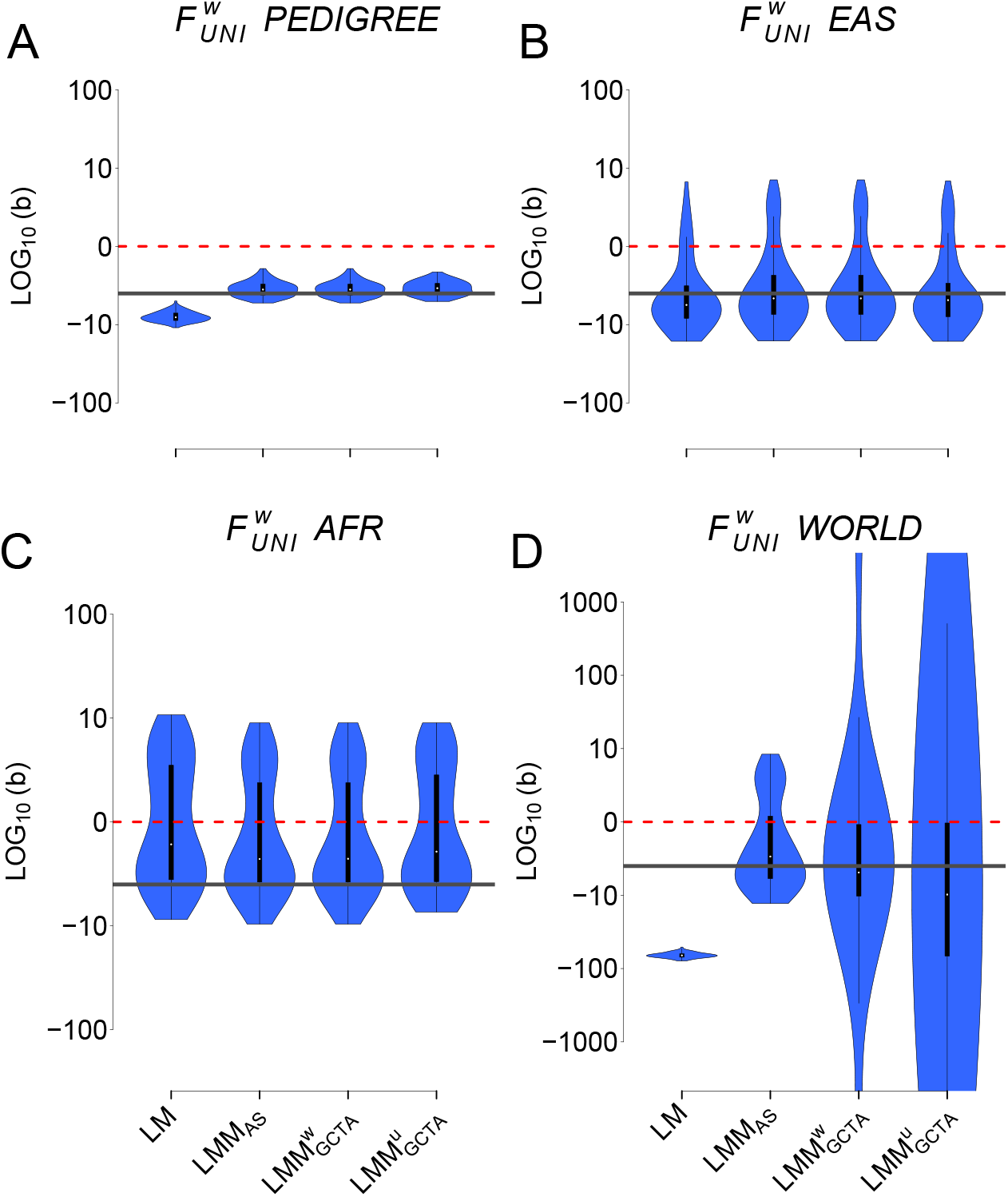
**Comparison of the inbreeding depression strength estimates (***b***) with** 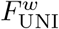 in the four populations with four different models. The four models are: i) the simple linear regression (LM), ii) the linear mixed model with the allele sharing relatedness matrix as a random factor, iii) the linear mixed model with the weighted `GCTA` relatedness matrix as a random factor and iv) the linear mixed model with the unweighted `GCTA` relatedness matrix as a random factor. Panel **A** the simulated PEDIGREE population, panel **B** depicts the EAS population, panel **C** the AFR population and finally panel **D** the WORLD population. Note that all panels are in *log*_10_ scale. Also note that LMM did not converge for some replicates (yielding estimated *b* values above 1000 or below - 1000, not shown in the graph). Percentages of replicates which did not converge: panel **D** (WORLD): 21% for 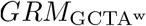; 20% for 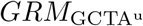.

### Distribution of additive and dominance effects

We found a difference between the three linear mixed models only because the scenario presented in the main text includes effect sizes and dominance coefficients proportional to causal markers’ MAF as well as DEMA. When none of these three parameters are included, there is little difference between the three linear mixed models (Figure S8, panels B, F, J, N vs panels C, G, K, O vs panels D, H, L, P; tables S2-S5). Additional simulations were conducted without additive and dominance coefficients proportional to loci’s MAF and DEMA to assess their impact on ID detection. The individual and pairwise effects of additive and dominance coefficients being proportional to MAF and DEMA (the other scenarios of table S1) are explored and discussed in details in supplementary material and Figures S8-S15.

Finally, we also investigated i) the effect of the LDMS stratification method proposed by Yengo [33] (Figures S8-S15) but found that it only improves results with the simple LM and ii) the effect of using intermediate frequencies causal loci (Figure S18) which reduced the variance in *b* estimates for all inbreeding coefficients.

## Discussion

By analyzing the phenotypes of a large simulated pedigreed polygamous population with high family structure as well as subsets of the 1000 genomes project [28], we demonstrated that, despite population or family structure, inbreeding depression estimates can be accurately measured if the data are analyzed with a mixed model using the genomic relationships among individuals as random effect. In comparison to the other genomic relationship matrices (GRMs), the one based on allele sharing provides the most consistent and accurate results, especially for smaller sample sizes and samples with a high family or population structure. And, among the several inbreeding estimators tested, 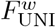 proved the most reliable to quantify inbreeding depression.

We observed trivial differences among the different models when there is no population structure (i.e. in the EAS and AFR populations). However, as soon as there is some structure (the WORLD and POLYPED populations) the classical linear model (LM) completely fails to estimate *b* regardless of the inbreeding coefficient used. This result is concordant with Yengo *et al*. (2017) [33] where the authors quantified ID using a simple linear model and demonstrated that *F*_HOM_ (whose properties are very similar to *F*_AS_), 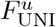 and two different *F*_ROH_ were sensitive to population structure. As for the comparison of three linear mixed models (LMM), they perform equally when there is no population structure (EAS and AFR) or very large sample sizes (11,924 individuals from the complete PEDIGREE population). Although samples of this size are common for research on humans, they will seldom be found in wild populations. We therefore subsampled the PEDIGREE population to 2,500 individuals in order to investigate the effect of a smaller sample size. We used two types of sub-sampling: i) random sub-sampling where individuals were chosen completely randomly and ii) ranged sub-sampling where individuals where chosen to maximise the range of *F* in the sampled population. We stress that what we consider a small sample size (2,500 individuals) will not be found in many wild species, particularly for endangered populations, where monitoring inbreeding and inbreeding depression are critical. As expected, when we sub-sampled individuals from the PEDIGREE population, RMSE values associated with *b* estimation increased slightly for both LMM_AS_ and 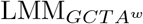 mixed models and we failed to detect ID in some replicates. Accordingly, even with 2,500 individuals, we lack power and several thousands of individuals would be required to detect ID efficiently as Keller et al. and Caballero et al. previously pointed out [16, 4]. With the 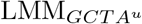 mixed model, all inbreeding coefficients but *F*_AS_ and *F*_UNI_ had convergence issues, suggesting that the 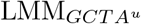 mixed model is the least robust of the three mixed models. As expected, randomly sub-sampling individuals lead to a larger variance of *b* estimates compared to the ranged sub-sampling scheme, indicating that maximizing the variance of samples’ *F* improves the estimation of *b*, although it is not obvious how such sampling could be done in non monitored natural populations. When we add strong population structure in addition to the small sample size (2,504 individuals from the highly structured WORLD population), we observe striking differences between the three different GRMs. The linear mixed model including the allele sharing based GRM (LMM_AS_) resulted in the most efficient estimations of *b*. In addition, the mixed models with both 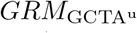 and 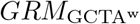 did not converge for high percentages of replicates (compared to 0% for LMM_AS_) emphasizing that LMM_AS_ is the best model for quantifying inbreeding depression in highly structured populations (although the most used GRM is currently the one estimated from `GCTA`). This is because the allele sharing based GRM matrix is a better estimator of kinship compared to both `GCTA` matrices [11, 30]. Indeed what the *GRM*_AS_ estimates is the actual kinship in the population, based on how many alleles individuals share. In contrast, what both 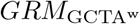 and 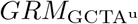 estimate is a combination of individual kinship, their mean kinship with the other individuals and the overal mean kinship in the population (see eq. 3 in Goudet *et al*. [30]). Consequently, since the kinship itself is better estimated with *GRM*_AS_, the non-independence of observations (and thus the population structure) is better accounted for with LMM_AS_ which leads to better *b* estimates. Importantly, the inclusion of a GRM in the ID estimation model is not limited to simple linear models. Even though we used only linear models in this study, any type of generalized linear model can incorporate a GRM as a random factor. Consequently this method can be applied to any trait distribution. Furthermore, by including the GRM-based random factor, the non-independence of observations is better accounted for than by including the population as a random factor, and no prior knowledge of the population structure is required.

### Comparing *F*

Concerning the different inbreeding coefficients, we found 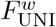 to be the best *F* for quantifying inbreeding depression. Indeed, 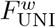 was the only coefficient we tested which was not sensitive to either additive and dominance effect sizes being proportional to MAF or DEMA resulting in the least biased estimation of *b*. On the contrary, we found that 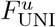 was influenced by the dominance effect sizes being proportional to MAF and by population structure. Since 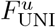 gives equal weight to all loci, the rare allele associated with large dominance coefficients add noise in the estimation of *b*. Similarly, when there is population structure, rare alleles which have strong influence on 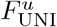 are likely to be private alleles which will strongly bias population-specific allelic frequencies and eventually 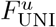 estimation. Importantly, 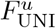 performed as well as 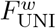 when we filtered on *MAF >* 0.05 for *F* and all GRMs estimation. This is because 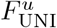 uses the average of ratios, which results in loci with small MAF strongly influencing the outcome. When these rare loci are filtered out, the estimated *F* is no longer biased. This explains why Yengo *et al*. [33] found that 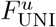 was the best *F* for quantifying inbreeding depression with an homogeneous subset of the UK bio bank dataset: they filtered on *MAF >* 0.05 leading to 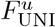 estimation not being influenced by rare alleles with strong additive and/or dominance effect sizes. Concerning *F*_AS_, we found that it was very sensitive to DEMA. This result is also concordant with Yengo *et al*. [33] who found that *F*_HOM_ (with properties very similar to *F*_AS_) was sensitive to DEMA. In this paper the authors explain that this sensitivity is due to *F*_HOM_ (and thus *F*_AS_) correlating strongly with minor allelic count which will create a spurious association with inbreeding depression in the presence of DEMA. However, *F*_AS_ resulted in the most accurate estimates of *b* when DEMA was not included in the model, suggesting that it is the best *F* to estimate inbreeding for neutral regions. Finally, we found that ROHs and HBD segments based *F*, namely *F*_ROH_ and *F*_HBD_, performed poorly: underestimating the strength of inbreeding depression (positive *b*) or displaying very large variance among replicates. This result is in contradiction with Kardos *et al*. [13, 14] and Nietlisbach *et al*. [25] who found that *F*_ROH_ and *F*_HBD_ were better at quantifying inbreeding depression compared to SNPs-independent based *F*. However, Alemu *et al*. [1] and Caballero *et al*. [5] showed the best *F* actually depends on the history of the population. Indeed, they showed that *F*_ROH_ and *F*_HBD_ and to a lesser extent *F*_HOM_ were better at quantifying homozygosity at loci with common alleles. On the contrary, 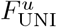 was better at quantifying homozygosity at rare alleles. Alemu *et al*. [1] and Caballero *et al*. [5] propose that, on the one hand, in populations with low effective sizes, selection is weaker and deleterious alleles may be able to reach intermediate frequencies as a result of drift. Therefore both *F*_ROH_ and *F*_HBD_ (and *F*_HOM_ in their analyses) should perform better in such populations. In our study, the standard scenario (with no ADD, no DOM and no DEMA) mimics what happens in such small populations and we found that *F*_ROH_, *F*_HBD_ and *F*_AS_ (which has similar properties to *F*_HOM_) performed better than 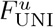 (which is the *F*_UNI_ they tested) in the highly structured WORLD population and to a lesser extent in the family structured PEDIGREE population. With homogeneous populations, we do not observe any difference between these inbreeding coefficients. Nevertheless, this is consistent with Alemu [1] results, as they used families which consequently create structure. On the other hand, in populations with a large effective size, selection maintains deleterious alleles at low frequencies which explains why Yengo *et al*. (2017) found that *F*_UNI_ was the best *F* with the large UK biobank dataset and this is consistent with what we have found with the ADD & DOM & DEMA scenario which mimics what happens in populations with large effective sizes.

## Conclusion

In this paper, we showed that the more accurate method for estimating inbreeding depression is to use a mixed model with an allele-sharing-based relatedness matrix as a random component but 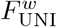 as the inbreeding coefficient to predict inbreeding depression. The most commonly used GRM 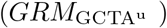 results in biased and highly variable estimates of *b* in structured populations. We stress that even if the results are greatly improved by using the allele-sharing GRM and 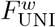, the variance among replicates is still large and no inbreeding depression is detected in several replicates 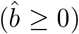 in the highly structured WORLD population as well as in the small and slightly admixed AFR population. Therefore, detecting efficiently inbreeding depression of the magnitude commonly found and that we simulated requires very large sample sizes with several thousand individuals, particularly in structured populations. Unfortunately, this might be hardly feasible for wild and/or endangered populations.

