## supplementary material for "Detecting inbreeding depression in structured populations"

#### Detailed introduction

Inbreeding depression, the decrease in mean phenotypic value in inbred individuals, is a phenomenon pervasive in humans, domestic and wild animals and plants [55, 26, 14]. Inbreeding has been associated with various diseases and is the result of mating between relatives. It has been observed in a wide range of taxa such as humans [3, 10], livestock [42, 29], wild animal populations [27] and plants [27, 57]. The effect of inbreeding on individuals' genomes is to increase homozygosity: since related individuals share higher genetic similarity, their offspring are likely to harbor higher fractions of homozygous identical-by-descent (IBD) genomic regions. Mating between closely related individuals such as siblings or first degree cousins results in strong inbreeding, also referred to as 'recent inbreeding'. For instance, the mating of closely related individuals is encouraged in domestic species, as part of the artificial selection of the process of domestication is mating individuals with similar phenotypes of economic interest [36, 41]. Nonetheless, recent inbreeding also occurs in wild isolated populations especially those with extremely small effective sizes [16, 13] and in various human populations where mating with relatives is culturally encouraged [4]. On the other hand, more moderate and remote inbreeding may occur because of ancient relatedness between the parents, which is often observed in populations with small effective sizes or ancient founder effect. This type of inbreeding has been observed in humans, notably in the individuals with East Asian ancestry from the 1,000 Genomes Project [18] but also happens in wild [48] and domestic [36] populations with small sizes.

#### What is Inbreeding Depression?

Inbreeding is often associated with reduced fitness, a phenomenon called inbreeding depression (ID), in many different species such as humans [9, 8], other animals [45, 21, 34], and plants [57]. Charlesworth and Willis suggested two specific mechanisms by which increased homozygosity leads to a reduction in population fitness [12]. The first mechanism is balancing selection: heterozygotes are at an advantage when there is balancing selection; inbred individuals have higher chances of being homozygous, thus they will tend to have lower fitness. This mechanism has been characterized in drosophila [12] but is likely to be of lesser importance in other species [12, 20]. The second mechanism involves partially recessive deleterious alleles which, when only present in one copy, have little effect and are therefore undetected by natural selection. Hence, selection can only act upon them when they are in the homozygous state. For partially recessive deleterious alleles, the strength of selection is weak, even in the homozygous state. As a result, many of these loci are segregating at low frequencies in populations [44]. For non-inbred individuals, the effect on fitness is minimal: they might carry some of these alleles in homozygous form by chance, but since most of these alleles are only partly detrimental, the individuals' overall fitness is hardly impacted. Conversely, the more inbred an individual is, the higher is its proportion of genome in the homozygous state, increasing the likelihood of a large quantity of partially deleterious alleles in a homozygous state. As these alleles accumulate, their deleterious effects will have a significant impact on the fitness of inbred individuals. In populations with small sizes, these marginally recessive alleles can easily reach intermediate frequencies [27] or even fixation [17] since the effect of drift will be much stronger. Moreover, individuals will tend to share more co-ancestry in these small populations, resulting in a large accumulation of these deleterious recessive alleles in the homozygous state.

#### Why measuring inbreeding?

Since it can have disastrous effects on populations, quantifying inbreeding and its deleterious consequences is of the utmost importance. In humans, for example, researchers were able to link inbreeding with many deleterious phenotypes [52, 43] which led to a better understanding of the underlying mechanisms involved in these traits. Quantifying inbreeding is also essential for monitoring endangered and small isolated populations. If their inbreeding status is high, we can expect a decline of the population in future generations. To avoid this issue, strict breeding programs which aim at reducing the overall inbreeding load of the population can be implemented [30, 47, 49].

#### How to quantify inbreeding ?

Many different methods have been developed for inbreeding quantification and there is no consensus on which one is the best [1, 7, 19, 40, 56, 58]. The classical approach was first proposed by Sewall Wright in 1922 and makes use of pedigrees (called hereafter $F_{\mathrm{PED}}$ ) [53]. $F_{\mathrm{PED}}$ corresponds to the probability that two alleles are IDB (a definition proposed by Malécot in 1948 [33]) and rely on the genealogy of the population. Consequently its estimation is only possible in populations where matings are actively recorded (i.e. mainly human and domestic populations). Furthermore, what $F_{\mathrm{PED}}$ measures is the expected inbreeding coefficient, which can be very different from the realized coefficient due to recombination stochasticity and random segregation of alleles. With the advances in sequencing technologies, genomic-based inbreeding coefficients (hereafter called $F_{\mathrm{genomic}}$ ) have been developed. Among these, some coefficients rely on the comparison between observed and expected heterozygosity such as $F_{\mathrm{HOM}}$ [11, 46], the expected allele sharing between individuals such as $F_{\mathrm{AS}}$ [58] or on the correlation between uniting gametes such as $F_{\mathrm{UNI}}$ [54]. In addition to estimating the realized inbreeding coefficient and requiring no prior knowledge of the mating behavior of the population, these genomic estimates are simple and straightforward to compute and do not require whole-genome sequencing (WGS) data; a few thousands SNPs are usually sufficient for reliable inbreeding estimation in humans [19]. However they also have a disadvantage: they usually rely on allelic frequencies (except for $F_{\mathrm{AS}}$ ) and therefore if these frequencies have not been correctly estimated, this will affect the estimation of these coefficients. Additionally, these coefficients treat each SNP independently, whereas in nature DNA is transmitted from parents to offspring in large chromosomal chunks. In order to take this into account, McQuillan *et al.* (2008) proposed a new inbreeding coefficient: $F_{\mathrm{ROH}}$ which uses runs of homozygosity (ROHs) long homozygous stretches as proxy for IBD segments within individuals [37]. A model-based approach relying on hidden Markov models has also been developed for detecting IBD segments [32] by identifying homozygous-by-descent (HBD) segments. This model is the basis for many other model-based IBD segments detection methods such as `BCFTools` [39], `BEAGLES` [5] and `RZooRoH` [15]. The inbreeding coefficient estimated with these model-based approaches will be called $F_{\mathrm{HBD}}$ from now on. One advantage of these methods is that they do not rely on allelic frequencies which can be very valuable when only a few individuals are available. However, it has been shown that these coefficients and especially $F_{\mathrm{ROH}}$ are sensitive to SNPs density and parameters sets and no consensus on what is the best set of parameters exists nowadays [38, 31].

### How to quantify inbreeding depression?

How to quantify inbreeding depression, although central to conservation genetics for decades [27] is still debated. This debate includes two sub-questions: which statistical model should be utilized ? And which inbreeding coefficient ? Regarding the model, the classical approach consisted of using linear regression of the phenotypes on the inbreeding coefficient. However, other models have been used, such as maximum likelihood and Generalized Linear models (GLMs) with various link functions. In 2019, Nietlisbach *et al.*. [40] compared different models and found that the common GLM models with logit link did not allow for accurate inbreeding depression strength estimation. They propose using maximum likelihood estimation or GLM with logarithm link functions.

#### Effect of sample size

Except for humans and domestic species where genetic and phenotypic data are available for several thousands of individuals [56, 51, 35, 45, 50], ID studies in the wild are usually performed on smaller sample sizes varying between 100 and few thousands individuals [21, 27, 25, 24]. Consequently, we may not be able to detect inbreeding depression in many wild populations (unless the effect is very strong). Indeed Keller et al. [28] stressed that to detect the effect of deleterious alleles with small effects (and $F_{\text{ROH}}$ ), very large sample sizes of thousands of individuals are needed.

#### Review of what have been done so far

An unresolved issue is which inbreeding coefficient is more accurate for quantifying inbreeding depression. In 2011, Keller *et al.* [28] performed simulations mimicking past human demography and compared different inbreeding coefficients. The authors showed that $F_{\text{ROH}}$ retains more individual variation compared to SNPs-independent measures of inbreeding and correlates best with homozygous mutation load which they suggest is likely to make it the best $F$ for quantifying inbreeding depression. However the authors stress that to detect the effect of deleterious alleles with small effects, very large sample sizes are needed. In 2015, Kardos *et al.* [22] also performed simulations and compared the ability of $F_{\text{PED}}$ , $F_{\text{ROH}}$ and $F_{\text{HOM}}$ to capture the true proportion of genome within IBD segments. They found similar results to [28]: $F_{\text{PED}}$ is outperformed by all $F_{\text{genomic}}$ and among the $F_{\text{genomic}}$ they tested, $F_{\text{ROH}}$ performed better. In 2016, Bérénos *et al.* [2] used pedigrees and several $F_{\text{genomic}}$ and showed that genomic based coefficients of inbreeding detect more inbreeding depression compared to $F_{\text{PED}}$ . In 2017, Yengo *et al.* [56] used an homogeneous subset of the UK biobank dataset (individuals of European ancestries exclusively, and with kinships less than 0.05) to simulate traits and compare various $F$. The results they found contradicted Kardos *et al.* [22]: they found $F_{\text{UNI}}$ to be the best coefficient to estimate ID and that $F_{\text{ROH}}$ -based estimates of ID tended to be overestimated and showed higher standard error. In addition, one aspect discussed in this paper but not elsewhere as far as we know, is the spurious effect that directional additive effect can have on estimates of inbreeding depression (what the authors called DEMA, for Directional Effect of Minor Alleles). For a trait linked to fitness, we expect most new mutations to have detrimental effects, diminishing the value of the trait. Selection will tend to remove these detrimental alleles, or maintain them at low frequencies. Many low frequency alleles would then be detrimental, leading to a negative DEMA. The authors showed that $F_{\text{HOM}}$ (and thus $F_{\text{AS}}$ since they have similar properties) is sensitive to DEMA while $F_{\text{UNI}}$ and $F_{\text{ROH}}$ are not. They also showed via simulations that all estimates of inbreeding depression are somewhat sensitive to population structure, $F_{\text{UNI}}$ being the least affected. They recommend estimating inbreeding using Linkage Disequilibrium (LD) score and Minor Allele Frequency (MAF) bins, and to sum the ID estimates from these bins as an overall estimate of ID for the trait. In response to this article, Kardos and coauthors [23] argued that $F_{\mathrm{UNI}}$ yielded better results than $F_{\mathrm{ROH}}$ because of to the method Yengo *et al.* (2017) [56] used to compare the performance of the different $F$s and that $F_{\mathrm{ROH}}$ is preferable for studying inbreeding depression. In 2019, Nietlisbach *et al.* [40] published a paper using simulations and compared the capacity of different $F$s to quantify ID. They used the inbreeding load as the gold standard and found that $F_{\mathrm{ROH}}$ was the coefficient which showed the highest correlation with inbreeding load. In 2020, Caballero *et al.* [7] used simulations and included several populations with different histories: they found that the best $F$ actually depends on the size of the population. $F_{\mathrm{ROH}}$ did a better job at quantifying ID in population with small effective size while $F_{\mathrm{UNI}}$ was better at predicting ID estimates in populations with large effective sizes. However, the authors stressed that the error in ID estimates were large in all situations. Finally, in 2021, Alemu *et al.* [1] used SNPs-array empirical cattle data for several groups of allelic frequencies and found that $F_{\mathrm{UNI}}$ and $F_{\mathrm{GRM}}$ are better at quantifying homozygosity at rare alleles while $F_{\mathrm{ROH}}$ and $F_{\mathrm{HOM}}$ are better for alleles at intermediate frequencies and correlate better with whole-genome homozygosity. Consequently, the authors suggest that the history of the population will play a key role in determining which $F$ is best to estimate ID. Indeed recessive deleterious alleles which should be those responsible for inbreeding depression are expected to segregate at low frequencies in large populations due to negative selection. On the contrary, in small populations, drift can increase these deleterious recessive alleles frequencies to reach intermediate frequencies which would make $F_{\mathrm{ROH}}$ and $F_{\mathrm{HOM}}$ better suited to detect ID.

#### Summary of what we did, found and propose

In this paper we simulated traits based on simulated as well as empirical WGS human data from populations with various sizes from the 1,000 Genomes project. We show that some $F$ are more sensitive to population structure and DEMA than others. We confirm only some of Yengo *et al.* [56] results. Importantly, we show that accounting for the non-independence of observations with a mixed model via an allele sharing based genomic relationship matrix (GRM) and using a modified version of $F_{\mathrm{UNI}}$ which gives more weight to common alleles resolves most of the issues raised by Yengo *et al.* [56].

### Supplementary Material and methods

#### Summary of the simulated scenarios

Table and figure S1

| Scenario | Additive effect sizes | Dominance coefficients | DEMA |
|---|---|---|---|
| Standard | Randomly assigned | Randomly assigned | No |
| ADD | Proportional to MAF | Randomly assigned | No |
| DOM | Randomly assigned | Proportional to MAF | No |
| DEMA | Randomly assigned | Randomly assigned | Yes |
| ADD & DOM | Proportional to MAF | Proportional to MAF | No |
| ADD & DEMA | Proportional to MAF | Randomly assigned | Yes |
| DOM & DEMA | Randomly assigned | Proportional to MAF | Yes |
| ADD & DOM & DEMA | Proportional to MAF | Proportional to MAF | Yes |

**Table S1: Simulated scenarios** used in this study. The first column corresponds to the scenario name and the three others indicate whether additive effect sizes (second column) and dominance coefficients (third column) were randomly assigned or according to MAF and whether DEMA (third column) was included.

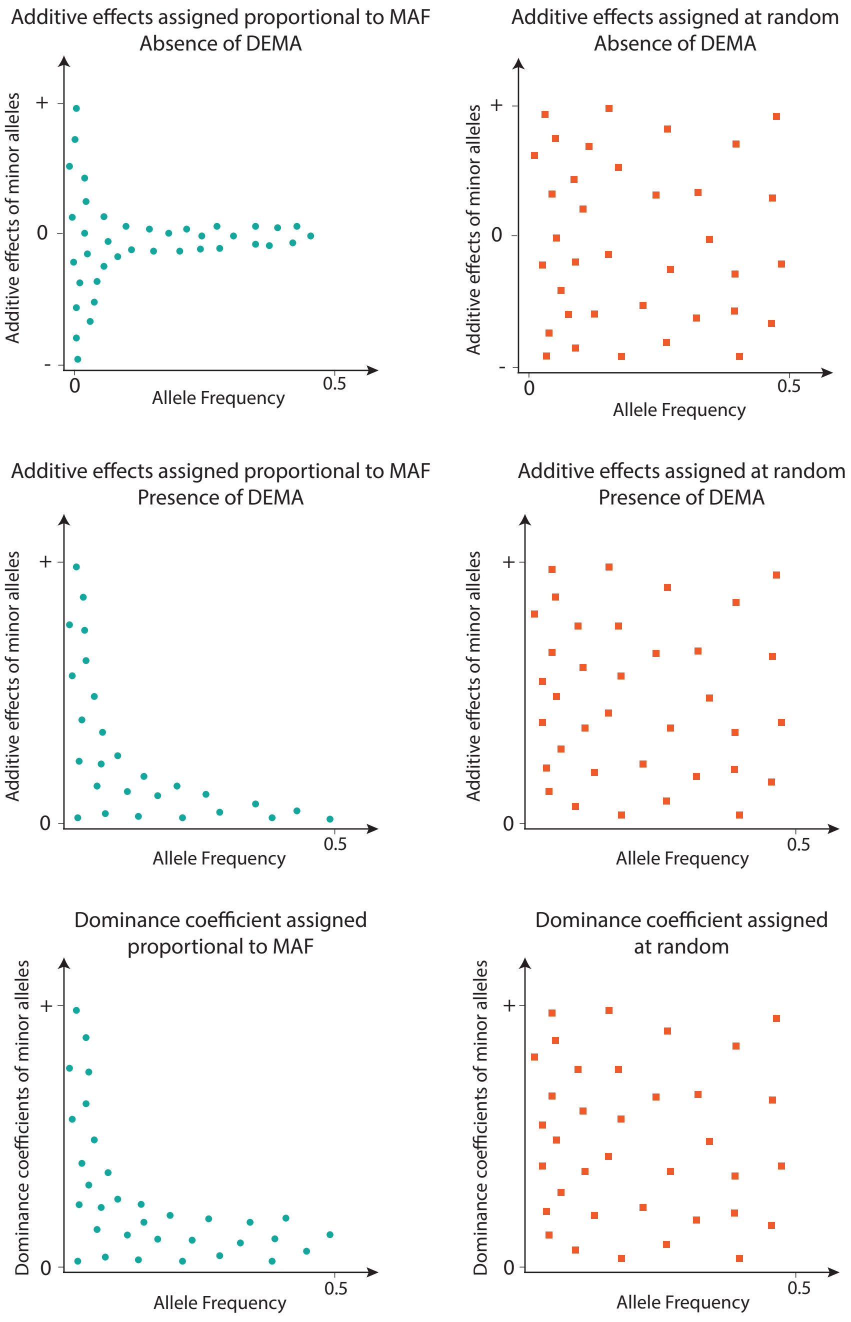


**Figure S1: Distribution of the additive effect sizes and dominance and dominance coefficients when proportional to MAF or randomly assigned in the absence and presence of DEMA.**

#### LDMS Stratification

For the SNPs-based inbreeding coefficients (i.e. $F_{\mathrm{AS}}$ and both $F_{\mathrm{UNI}}$ ), we also investigated whether using the Linkage disequilibrium and minor allele frequency stratified inference (hereafter called LDMS stratification) proposed by Yengo *et al.* (2017) [56] improved the estimation of the inbreeding depression strength ($b$). For LDMS stratification, each $F$ is estimated from each combination of 7 MAF bins and 4 LD bins, and the ID estimate is obtained as the sum of the partial regression coefficients of the trait on each of the 28 inbreeding coefficients. The MAF and LD bins are defined as in [56]: $MAF_1 : p \leq 0.001$; $MAF_2 : 0.001 < p \leq 0.01$; $MAF_3 : 0.01 < p \leq 0.1$; $MAF_4 : 0.1 < p \leq 0.2$; $MAF_5 : 0.2 < p \leq 0.3$; $MAF_6 : 0.3 < p \leq 0.4$; $MAF_7 : 0.4 < p \leq 0.5$, and LD bins correspond to the 4 quartiles. For EAS samples, since there is only 500 samples, we discarded $MAF_1$ and made $MAF_2 : p \leq 0.01$. Results for this analyses can be found in figures S8-S15.

#### MAF filtering

In order to verify whether rare alleles were responsible for $F^u_{\mathrm{UNI}}$ poor estimations of $b$, we did as Yengo *et al.* (2017) [56] and filtered the WORLD populations genomic data to keep only SNPs with MAF $> 0.05$ using `BCFTools`. We then re-estimated all $F$ and GRMs on the newly filtered data set and re-simulated inbreeding depression. Results for this analysis can be found in figure S16.

#### HBD segments without size selection

When estimating IBD segments in the genome, an advantage of model-based approaches (such as `BCFTools`) is that there is no constraint on the minimum size of an HBD segment. However, since the coalescing events responsible for inbreeding depression are usually recent, we only considered segments larger than 100KB and 1MB in the main text. We wanted to test whether using all HBD segments independently of their size would yield better $b$ estimates. Consequently, in the WORLD population and from the output of `BCFTools`, we did not filter on size but rather on quality score. Indeed, the `BCFTools` output includes a quality score which gives an indication about how confident we are about an HBD segment being IBD. A minimum quality of 30 was used for filtering. Results for this analysis can be found in figure S17.

### Supplementary Results and Discussion

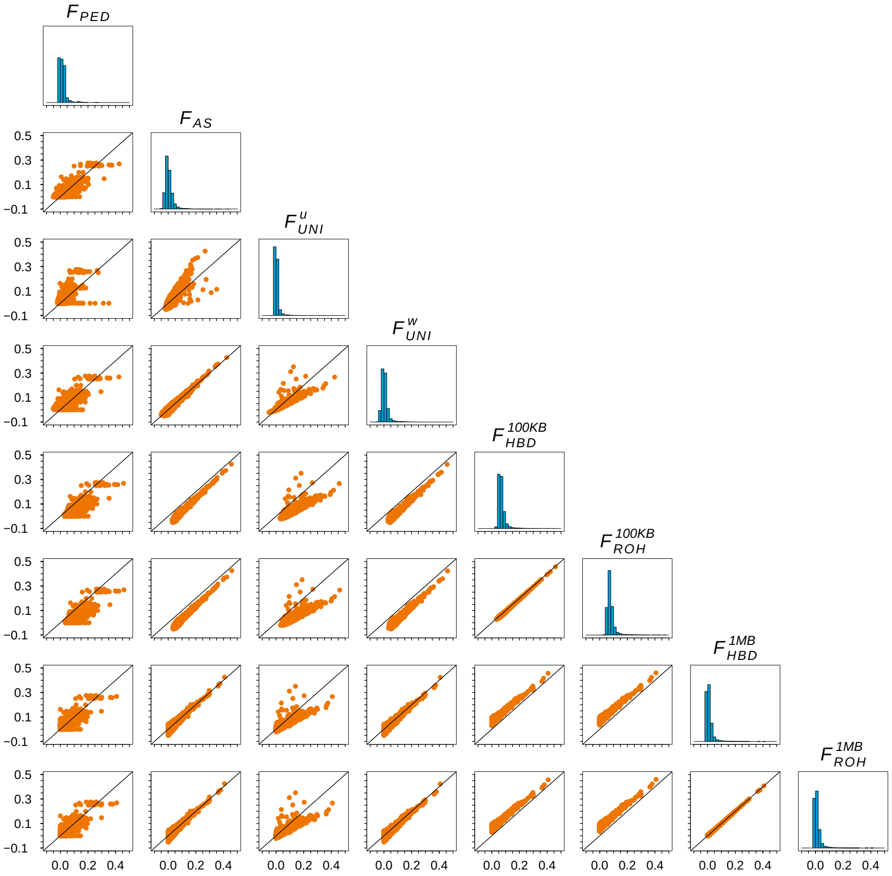


**Figure S2: pairwise comparison among the different inbreeding estimates ($F$) in the simulated PEDIGREE population.** $F$ depicted in this figure are $F_{\text{PED}}$ , $F_{\text{AS}}$ , $F^{u}_{\text{UNI}}$ , $F^{w}_{\text{UNI}}$ , $F^{100KB}_{\text{HBD}}$ , $F^{100KB}_{\text{ROH}}$ , $F^{1MB}_{\text{HBD}}$ and $F^{1MB}_{\text{ROH}}$ .

Figure S2 shows the comparison among the different inbreeding coefficients used in the PEDIGREE population. In general, we can see that there is a strong correlation between all $F_{\text{genomic}}$ , with the exception of $F^{u}_{\text{UNI}}$ and the other $F$. In addition, the non-genomic-based $F_{\text{PED}}$ has the lowest correlation with all the other $F$.

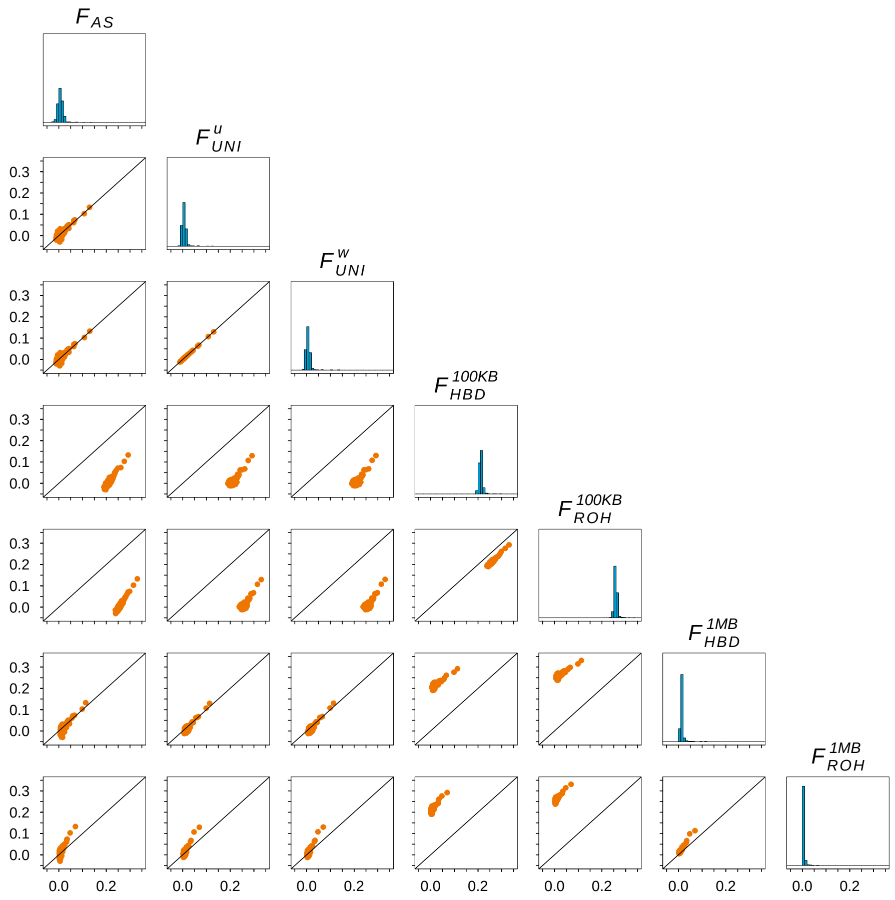


**Figure S3: pairwise comparison among the different inbreeding estimates ($F$) in the 1,000 Genomes Project EAS population.** $F$ depicted in this figure are $F_{\text{AS}}$ , $F^{u}_{\text{UNI}}$ , $F^{w}_{\text{UNI}}$ , $F^{100KB}_{\text{HBD}}$ , $F^{100KB}_{\text{ROH}}$ , $F^{1MB}_{\text{HBD}}$ and $F^{1MB}_{\text{ROH}}$ .

Figure S3 shows the comparison among the different inbreeding coefficients used in the EAS population. With no structure in the population, there is a good correlation between all $F_{\text{genomic}}$ . Even though the absolute values of the $F$ are different, the rank of inbreeding is always conserved among individuals (i.e. the most inbred individuals are the most inbred for all $F$).

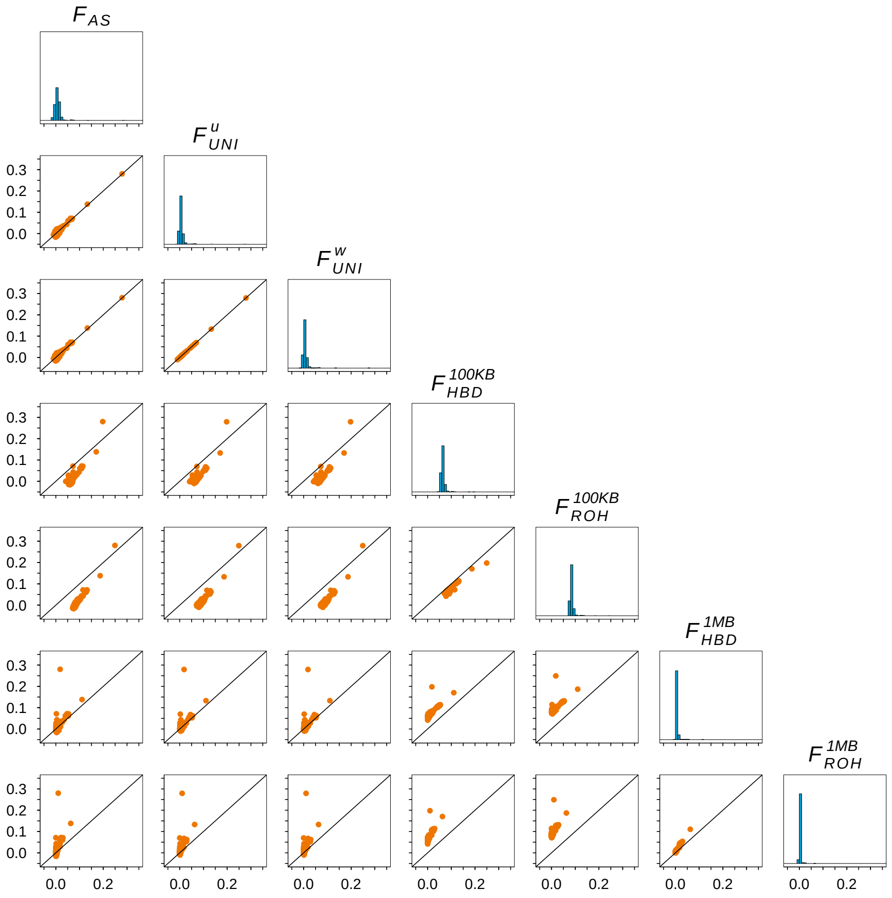


**Figure S4: pairwise comparison among the different inbreeding estimates ($F$) in the 1,000 Genomes Project AFR population.** $F$ depicted in this figure are $F_{\text{AS}}$ , $F^{u}_{\text{UNI}}$ , $F^{w}_{\text{UNI}}$ , $F^{100KB}_{\text{HBD}}$ , $F^{100KB}_{\text{ROH}}$ , $F^{1MB}_{\text{HBD}}$ and $F^{1MB}_{\text{ROH}}$ .

Figure S4 shows the comparison among the different inbreeding coefficients used in the AFR population. In this population too, there is a good correlation between all $F_{\text{genomic}}$ (especially the three SNPs-based $F$: $F_{\text{AS}}$ , $F^{u}_{\text{UNI}}$ and $F^{w}_{\text{UNI}}$ ).

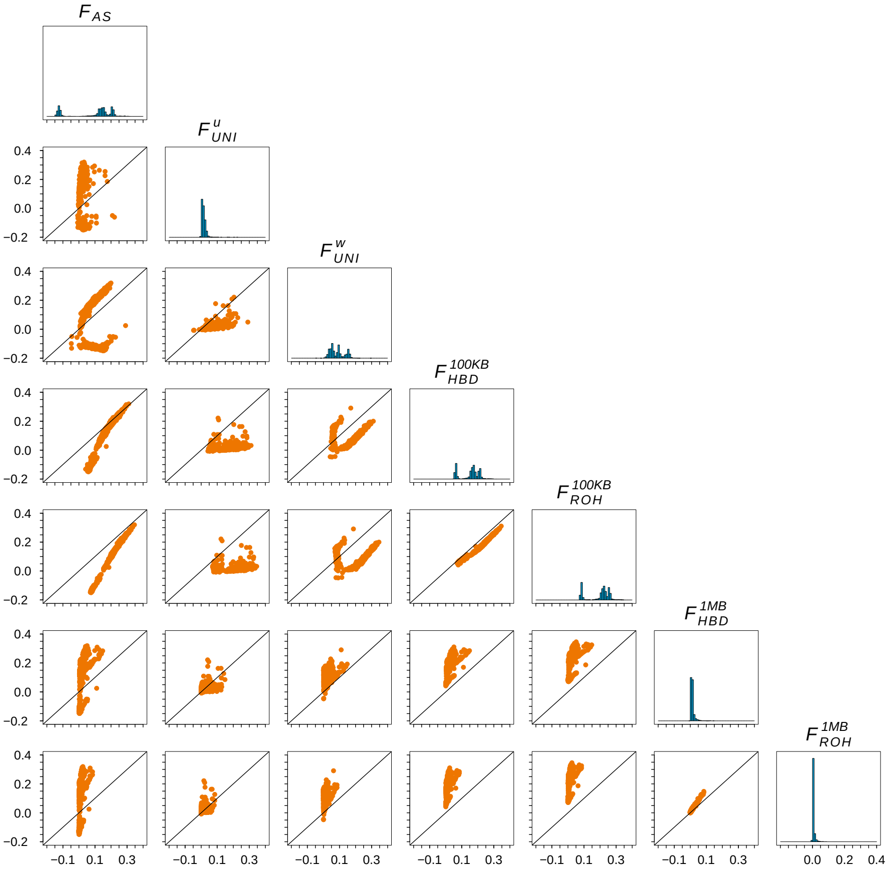


**Figure S5: pairwise comparison among the different inbreeding estimates ($F$) in the 1,000 Genomes Project WORLD population (all the individuals).** $F$ depicted in this figure are $F_{\mathrm{AS}}$ , $F^{u}_{\mathrm{UNI}}$ , $F^{w}_{\mathrm{UNI}}$ , $F^{100KB}_{\mathrm{HBD}}$ , $F^{100KB}_{\mathrm{ROH}}$ , $F^{1MB}_{\mathrm{HBD}}$ and $F^{1MB}_{\mathrm{ROH}}$ .

Figure S5 shows the comparison among the different inbreeding coefficients used in the structured WORLD population. In this population as well, there is a good correlation between all $F_{\mathrm{genomic}}$ except $F^{u}_{\mathrm{UNI}}$ and the others. In addition, among the other $F$, the African samples are the ones with the lowest correlation.

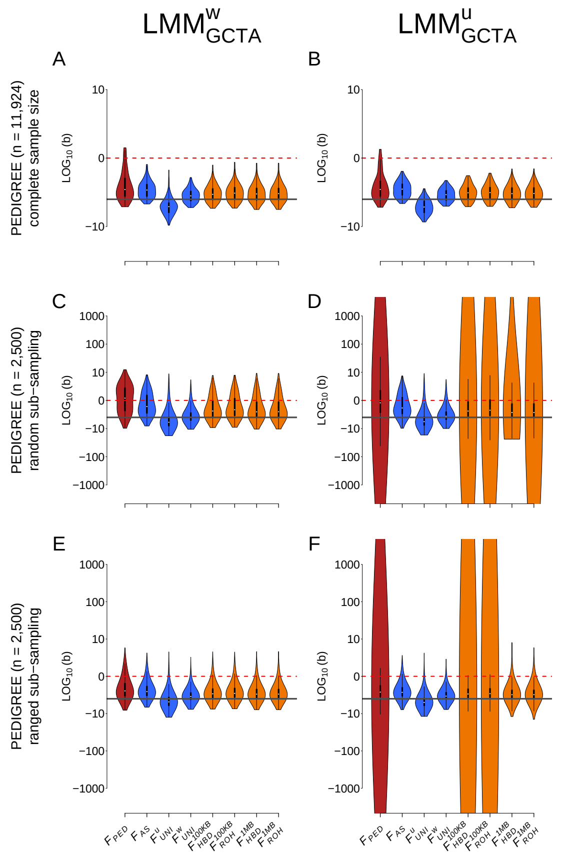


**Figure S6: Comparison of the estimation of inbreeding depression strength ($b$)** among different $F$ estimates and the LMM including two different GRMs in the PEDIGREE population and with the ADD & DOM & DEMA scenario. The first column depicts the LMM including the $GCTA^w$ matrix (panels **A**, **C** and **E**) and the second column the linear mixed model with the $GCTA^u$ matrix (panels **B**, **D** and **F**). The first row shows the complete simulated population (n = 11,924 individuals) on panels **A** and **B**. The second row depicts the randomly sub-sampled population (n = 2,500 individuals) on panels **C** and **D** and the third row shows the ranged sub-sampled PEDIGREE population (n = 2,500 individuals) on panels **E** and **F**. Inbreeding estimates presented in this graph are $F_{\text{PED}}$ , $F_{\text{AS}}$ , $F^{u}_{\text{UNI}}$ , $F^{w}_{\text{UNI}}$ , $F^{100KB}_{\text{HBD}}$ , $F^{100KB}_{\text{ROH}}$ , $F^{1MB}_{\text{HBD}}$ and finally $F^{1MB}_{\text{ROH}}$ . For panels **A** and **B**, violin plots represent the distribution of the inbreeding depression strength estimates ($b$) among the simulated 100 replicates. For panels **C** to **F**, violin plots represent the distribution of the inbreeding depression strength estimates ($b$) among the 10,000 simulated and sub-sampling replicates (100 sub-sampling replicate for each of the 100 simulation replicates). The solid dark grey line is the true strength of ID ($b$ = -3). The dashed red line represents the absence of ID ($b$ = 0), meaning that we failed to detect ID in any replicate above this line. Note that all panels are in log10 scale.

Figure S6 presents the inbreeding depression (ID) strength estimates ($b$) for the different inbreeding coefficients ($F$), with two models in the PEDIGREE populations and with the ADD & DOM & DEMA scenario. The first and second columns depict $b$ estimated with LMM using the unweighted ($LMM_{GCTA^u}$ ) and weighed ($LMM_{GCTA^w}$ ) `GCTA` matrices as random factors, respectively. The first row shows results for the complete PEDIGREE population (n = 11,924). The second row shows results for a reduced sample size of the PEDIGREE population (n = 2,500, meant to match the size of the 1KG WORLD population) where sub-sampled individuals were chosen completely randomly. The third row also shows results for a reduced sample size of the PEDIGREE population (n = 2,500) but these individuals were selected to represent the entire spectrum of inbreeding statuses. The violin plots show $b$ estimates distributions among the simulation replicates (100 replicates for the complete population, 10,000 replicates for both sub-sampled populations). The solid dark grey line is the true strength of ID ($b$ = -3). The dashed red line represents the absence of ID ($b$ = 0), indicating that we failed to detect ID in any replicate above this line. Root mean square error (RMSE) values associated with both regression models and populations are shown in main table 1. In the complete PEDIGREE population, we see little difference between the three GRMs we tested (figure 1, panel B VS figure S6, panels A and B; table 1): all $F$ yielded accurate estimates of $b$ when used inside a $lMM$, except for $F^u_{\text{UNI}}$ that slightly overestimates the strength of ID while $F_{\text{PED}}$ slightly underestimates it. However, when the sample size is reduced to 2,500 individuals, the strength of ID cannot be correctly estimated with $F_{\text{PED}}$ , $F^{100KB}_{\text{ROH}}$ and $F^{100KB}_{\text{HBD}}$ and the $LMM_{GCTA^u}$ model. In addition only the ranged sub-sampling allowed correct estimation of $b$ with both $F^{1MB}_{\text{ROH}}$ and $F^{1MB}_{\text{HBD}}$ and the $LMM_{GCTA^u}$ model. This confirms, first than $LMM_{GCTA^u}$ is the least robust among the regression models we used, and second that $F_{\text{ROH}}$ and $F_{\text{HBD}}$ based on larger segments are better when studying inbreeding depression.

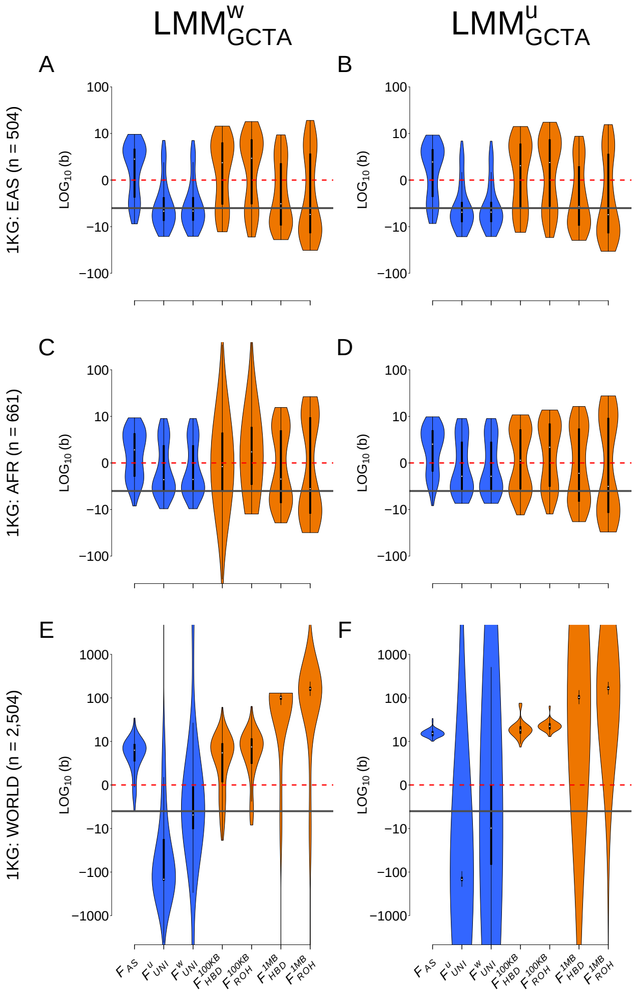


**Figure S7: Comparison of the estimation of inbreeding depression strength** ($b$) among different $F$ estimates and the $LMM$ including two different GRMs in the three populations from the 1,000 Genomes Project dataset and with the ADD & DOM & DEMA scenario. The first column depicts the LMM including the $GCTA^w$ matrix (panels **A**, **C** and **E**) and the second column the linear mixed model with the $GCTA^u$ matrix (panels **B**, **D** and **F**). The three rows show the three populations from the 1,000 Genomes project: EAS on panels **A** and **B**, AFR on panels **C** and **D** and WORLD on panels **E** and **F**. Inbreeding estimates presented are $F_{\text{AS}}$ , $F^{u}_{\text{UNI}}$ , $F^{w}_{\text{UNI}}$ , $F^{100KB}_{\text{HBD}}$ , $F^{100KB}_{\text{ROH}}$ , $F^{1MB}_{\text{HBD}}$ and finally $F^{1MB}_{\text{ROH}}$ . Violin plots represent the distribution of the inbreeding depression strength estimates ($b$) among the 100 simulations replicates. The solid dark grey line is the true strength of ID ($b$ = -3). The dashed red line represents the absence of ID ($b$ = 0), meaning that we failed to detect ID in any replicate above this line. Note that all panels are in $log_{10}$ scale.

Figure S7 presents the inbreeding depression (ID) strength estimates ($b$) for the different inbreeding coefficients ($F$), with two models in the three populations from the 1,000 Genomes Project: EAS, AFR and WORLD and with the ADD & DOM & DEMA scenario. The first and second columns show $b$ estimated with LMM respectively including the unweighted ($LMM_{GCTA^u}$) and weighed ($LMM_{GCTA^w}$) `GCTA` matrices as random factors. The first row shows results for the EAS population (n = 504), the second row shows results the AFR population (n = 661) and the third row shows results for the complete WORLD population (n = 2,504). The violin plots show $b$ estimates distributions among the simulation replicates (100 replicates). The solid dark grey line is the true strength of ID ($b$ = -3). The dashed red line represents the absence of ID ($b$ = 0), indicating that we failed to detect ID in any replicate above this line. RMSE values associated with both regression models and the three populations are shown in main table 2. In the EAS homogeneous population, we see little differences among the three mixed models (figure 2, panel B VS figure S7, panels A and B; table 2). There is also little difference between the three LMM for the three SNPs-based $F$ (i.e. $F_{\text{AS}}$ , $F_{\text{UNI}}^u$ and $F_{\text{UNI}}^w$ ) in the AFR population (panels C and D). However $F_{\text{ROH}}$ and $F_{\text{HBD}}$ including smaller segments resulted in larger variance among $b$ estimates with the $LMM_{GCTA^w}$ model (panel C). This was not the case for both $LMM_{\text{AS}}$ and $LMM_{GCTA^u}$ models. The variance among $b$ estimates was larger for the AFR population compared to the EAS population. This might be explained by the fact that almost all individuals in the AFR population have a $F$ close to 0. In the EAS population however, the variance in $F$ is larger. The larger RMSE value may also be due to admixture in some AFR individuals for whom $F$ estimation is more complex.

Finally, none of the `GCTA`-based GRM yielded accurate estimation of $b$ in the highly structured WORLD population. For this last reason, we select the allele-sharing GRM as the best GRM for estimating inbreeding depression.

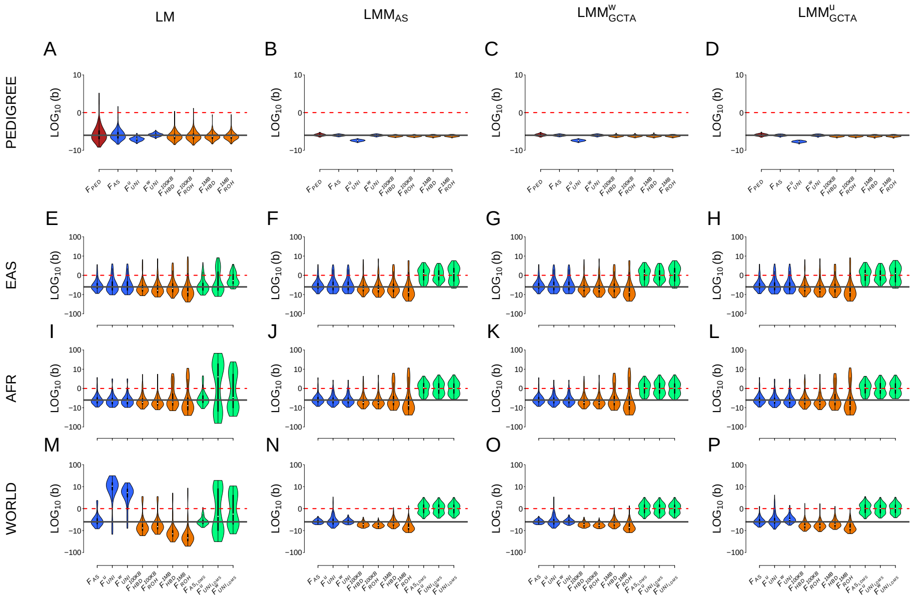


**Figure S8: Comparison of the estimation of inbreeding depression strength ($b$) among different $F$ estimates and models in four different populations with the standard scenario: effect sizes and dominance coefficients randomly assigned to each causal marker and no DEMA.** Each column represents a regression model. The first column depicts the simple linear regression (panel A, E, I and M), the second column the linear mixed model with allele sharing GRM matrix as random factor (panel B, F, J and N), the third column the linear mixed model with the $GCTA^w$ relatedness matrix as random factor (panel C, G, K and O) and finally the forth column represents the linear mixed model with $GCTA^u$ relatedness matrix as random factor (panel D, H, L and P). The first row depicts the complete simulated population (11,924 individuals): PEDIGREE in panels A, B, C and D. Inbreeding estimates compared in these panels (A - D) are $F_{\text{PED}}$ , $F_{\text{AS}}$ , $F^{u}_{\text{UNI}}$ , $F^{w}_{\text{UNI}}$ , $F^{100KB}_{\text{HBD}}$ , $F^{100KB}_{\text{ROH}}$ , $F^{1MB}_{\text{HBD}}$ and $F^{1MB}_{\text{ROH}}$ .The last three rows are the populations from the 1,000 Genomes Project: EAS in panels E, F, G and H, AFR in panels I, J, K and L and WORLD in panels M, N, O and P. Inbreeding estimates compared in these panels (E - P) are $F_{\text{AS}}$ , $F^{u}_{\text{UNI}}$ , $F^{w}_{\text{UNI}}$ , $F^{100KB}_{\text{HBD}}$ , $F^{100KB}_{\text{ROH}}$ , $F^{1MB}_{\text{HBD}}$ , $F^{1MB}_{\text{ROH}}$ , $F_{\text{AS}_{\text{LDMS}}}$ , $F^{u}_{\text{UNI}_{\text{LDMS}}}$ and finally $F^{w}_{\text{UNI}_{\text{LDMS}}}$ . Violin plots represent the distribution of the inbreeding depression strength estimates (b) among the 100 replicates. The solid dark grey line is the true strength of ID ($b$ = -3). The dashed red line represents the absence of ID ($b = 0$), meaning that we failed to detect ID in any replicate above this line. Note that all panels (A - P) are in $log_{10}$ scale.

Figure S8 presents the results of ID strength estimation for the standard scenario (additive effect sizes and dominance coefficients are randomly drawn and there is no DEMA). Corresponding RMSE values can be found in tables S2-S5. In the complete pedigree population, we can see that all regression models (but especially the mixed models) and all inbreeding coefficients result in efficient estimates (we use efficient to describe an estimate with low RMSE, thus which is unbiased and has low variance) (panels A-D, tables S3-S5). The most efficient estimate was $F^{w}_{\mathrm{UNI}}$ . The variance among $b$ estimates was larger for the three populations of the 1,000 Genomes Project and some replicates resulted in estimated $b$ above 0 (panels E-P). This is due to the smaller sample sizes. In both the EAS and AFR populations, $F_{\mathrm{AS}}$ , $F_{\mathrm{UNI}}$ and $F_{\mathrm{ROH}}$ gave unbiased estimates for all models (panels E-L). However, LDMS-based $F$ were always biased, especially for all the mixed models where they were centered around 0 (panels E-L, tables S3-S5). We believe it is because the sample sizes we used were too small to correctly estimate allelic frequencies per MAF and LD bins. With this scenario, $F_{\mathrm{AS}}$ (and $F_{\mathrm{AS}_{\mathrm{LDMS}}}$ in the simple LM model) gives the most efficient estimate of ID in the WORLD population (panels E-P, tables S3-S5). In addition, the three LMM perform similarly.

To conclude, when additive and dominance effects are uniform, and there is no DEMA, $F_{\mathrm{AS}}$ results in efficient estimates of ID (even with a simple linear model and strong population structure: panel M). With a mixed model, it is possible to estimate ID correctly for all inbreeding coefficients, except for LDMS-based $F$.

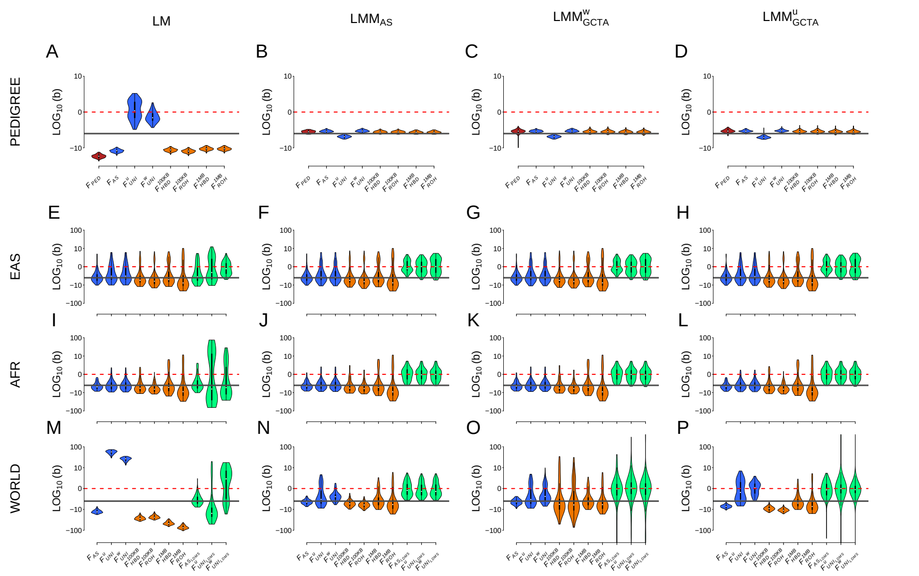


**Figure S9: Comparison of the estimation of inbreeding depression strength ($b$) among different $F$ estimates and models in four different populations with the ADD scenario: effect sizes assigned proportional to the MAF of causal markers, dominance coefficients randomly assigned to causal markers and no DEMA.** Each column represents a regression model. The first column depicts the simple linear regression (panel A, E, I and M), the second column the linear mixed model with allele sharing GRM matrix as random factor (panel B, F, J and N), the third column the linear mixed model with the $GCTA^w$ relatedness matrix as random factor (panel C, G, K and O) and finally the forth column represents the linear mixed model with $GCTA^u$ relatedness matrix as random factor (panel D, H, L and P). The first row depicts the complete simulated population (11,924 individuals): PEDIGREE in panels A, B, C and D. Inbreeding estimates compared in these panels (A - D) are $F_{\text{PED}}$ , $F_{\text{AS}}$ , $F^u_{\text{UNI}}$ , $F^w_{\text{UNI}}$ , $F^{100KB}_{\text{HBD}}$ , $F^{100KB}_{\text{ROH}}$ , $F^{1MB}_{\text{HBD}}$ and $F^{1MB}_{\text{ROH}}$ .The last three rows are the populations from the 1,000 Genomes Project: EAS in panels E, F, G and H, AFR in panels I, J, K and L and WORLD in panels M, N, O and P. Inbreeding estimates compared in these panels (E - P) are $F_{\text{AS}}$ , $F^u_{\text{UNI}}$ , $F^w_{\text{UNI}}$ , $F^{100KB}_{\text{HBD}}$ , $F^{100KB}_{\text{ROH}}$ , $F^{1MB}_{\text{HBD}}$ , $F^{1MB}_{\text{ROH}}$ , $F_{\text{AS}_{\text{LDMS}}}$ , $F^u_{\text{UNI}_{\text{LDMS}}}$ and finally $F^w_{\text{UNI}_{\text{LDMS}}}$ . Violin plots represent the distribution of the inbreeding depression strength estimates (b) among the 100 replicates. The solid dark grey line is the true strength of ID ($b$ = -3). The dashed red line represents the absence of ID ($b$ = 0), meaning that we failed to detect ID in any replicate above this line. Note that all panels (A - P) are in $log_{10}$ scale. Also note that linear mixed models did not converge for some replicates (yielding estimated $b$ values above 1000 or below -1000, not shown if outside the graph limits). **Percentages of replicates which did not converge:** panel O (WORLD, $GCTA^w$ ): 1% for $F_{\text{AS}_{\text{LDMS}}}$ , 3% for $F^u_{\text{UNI}_{\text{LDMS}}}$ and 1% for $F^w_{\text{UNI}_{\text{LDMS}}}$ ; panel P (WORLD, $GCTA^u$ ): 3% for $F^u_{\text{UNI}_{\text{LDMS}}}$ and 2% for $F^w_{\text{UNI}_{\text{LDMS}}}$ .

Figure S9 presents the results of ID strength estimation for the ADD scenario (when the additive effects are inversely proportional to MAF, the dominance effects are independent of MAF and there is no DEMA). Corresponding RMSE values can be found in tables S2-S5. The simple LM results in biased $b$ estimates with an overestimation of ID for all inbreeding coefficients except both $F_{\mathrm{UNI}}$ , which underestimate it (panel A, tables S3-S5). The three mixed models, on the other hand, provide efficient estimates of inbreeding depression in the PEDIGREE population (panels B-D, tables S3-S5). In the EAS population, all but the LDMS-based $F$ are unbiased, but with a large variance and we see no improvement of using a LMM rather than a simple LM (panels E-H, tables S3-S5). Interestingly, we see similar results for the AFR population but with lower variance (panels I-L, tables S3-S5). All estimates in the WORLD population, however, are strongly biased with the simple LM: ID is underestimated with $F_{\mathrm{AS}}$ and all $F_{\mathrm{ROH}}$ and overestimated with both $F_{\mathrm{UNI}}$ (panel M, tables S3-S5). For the WORLD population and the three LMM, $F_{\mathrm{AS}}$ yields the most efficient estimation of $b$, especially in combination with the $LMM_{\mathrm{AS}}$ model (for which the variance around $b$ estimates is the smallest) (panels N-P, tables S3-S5).

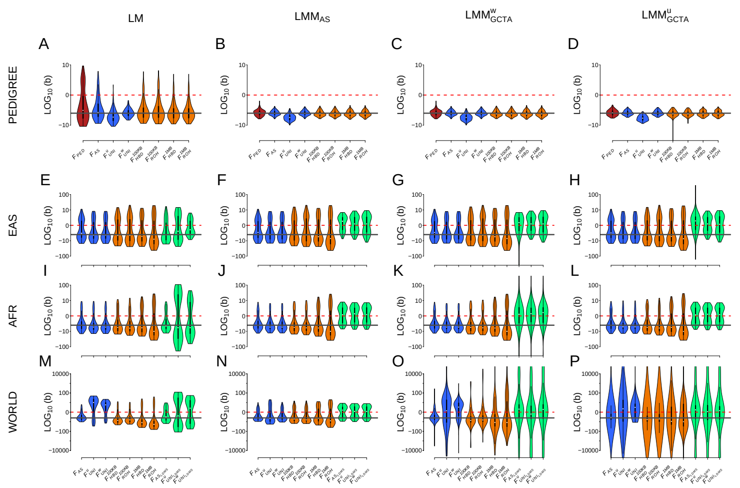


**Figure S10: Comparison of the estimation of inbreeding depression strength ($b$) among different $F$ estimates and models in four different populations with the DOM scenario: effect sizes randomly assigned to causal markers, dominance coefficients assigned proportional to the MAF of causal markers and no DEMA.** Each column represents a regression model. The first column depicts the simple linear regression (panel A, E, I and M), the second column the linear mixed model with allele sharing GRM matrix as random factor (panel B, F, J and N), the third column the linear mixed model with the $GCTA^w$ relatedness matrix as random factor (panel C, G, K and O) and finally the forth column represents the linear mixed model with $GCTA^u$ relatedness matrix as random factor (panel D, H, L and P). The first row depicts the complete simulated population (11,924 individuals): PEDIGREE in panels A, B, C and D. Inbreeding estimates compared in these panels (A - D) are $F_{\mathrm{PED}}$ , $F_{\mathrm{AS}}$ , $F^u_{\mathrm{UNI}}$ , $F^w_{\mathrm{UNI}}$ , $F^{100KB}_{\mathrm{HBD}}$ , $F^{100KB}_{\mathrm{ROH}}$ , $F^{1MB}_{\mathrm{HBD}}$ and $F^{1MB}_{\mathrm{ROH}}$ .The last three rows are the populations from the 1,000 Genomes Project: EAS in panels E, F, G and H, AFR in panels I, J, K and L and WORLD in panels M, N, O and P. Inbreeding estimates compared in these panels (E - P) are $F_{\mathrm{AS}}$ , $F^u_{\mathrm{UNI}}$ , $F^w_{\mathrm{UNI}}$ , $F^{100KB}_{\mathrm{HBD}}$ , $F^{100KB}_{\mathrm{ROH}}$ , $F^{1MB}_{\mathrm{HBD}}$ , $F^{1MB}_{\mathrm{ROH}}$ , $F_{\mathrm{AS_{LDMS}}}$ , $F^u_{\mathrm{UNI_{LDMS}}}$ and finally $F^w_{\mathrm{UNI_{LDMS}}}$ . Violin plots represent the distribution of the inbreeding depression strength estimates (b) among the 100 replicates. The solid dark grey line is the true strength of ID ($b$ = -3). The dashed red line represents the absence of ID ($b$ = 0), meaning that we failed to detect ID in any replicate above this line. Note that all panels (A - P) are in $log_{10}$ scale. Also note that linear mixed models did not converge for some replicates (yielding estimated b values above 1000 or below -1000, not shown if outside the graph limits). **Percentages of replicates which did not converge:** panel K (AFR, $GCTA^w$ ): 8% for $F_{\mathrm{AS_{LDMS}}}$ , 7% for $F^u_{\mathrm{UNI_{LDMS}}}$ and 4% for $F^w_{\mathrm{UNI_{LDMS}}}$ ; panel O (WORLD, $GCTA^w$ ): 1% for $F_{\mathrm{AS}}$ , 10% for $F^u_{\mathrm{UNI}}$ , 1% for $F^w_{\mathrm{UNI}}$ , 6% for $F^{100KB}_{\mathrm{HBD}}$ , 7% for $F^{100KB}_{\mathrm{ROH}}$ , 8% for $F^{1MB}_{\mathrm{HBD}}$ , 5% for $F^{1MB}_{\mathrm{ROH}}$ , 22% for $F_{\mathrm{AS_{LDMS}}}$ , 31% for $F^u_{\mathrm{UNI_{LDMS}}}$ and 31% for $F^w_{\mathrm{UNI_{LDMS}}}$ ; panel P (WORLD, $GCTA^u$ ): 10% for $F_{\mathrm{AS}}$ , 9% for $F^u_{\mathrm{UNI}}$ , 2% for $F^w_{\mathrm{UNI}}$ , 7% for $F^{100KB}_{\mathrm{HBD}}$ , 7% for $F^{100KB}_{\mathrm{ROH}}$ , 5% for $F^{1MB}_{\mathrm{HBD}}$ , 2% for $F^{1MB}_{\mathrm{ROH}}$ , 22% for $F_{\mathrm{AS_{LDMS}}}$ , 31% for $F^u_{\mathrm{UNI_{LDMS}}}$ and 28% for $F^w_{\mathrm{UNI_{LDMS}}}$ .

Figure S10 shows the DOM scenario (where dominance effects are inversely proportional to MAF while additive effects are independent and there is no DEMA). Corresponding RMSE values can be found in tables S2-S5. In the PEDIGREE population and with the simple LM, only $F^{w}_{\text{UNI}}$ gives efficient estimation of $b$; the other inbreeding coefficients are unbiased, but with large variance (panel A, table S2). For the same PEDIGREE population, the three mixed models reduce the variance around $b$ estimates (panels B-D, tables S3-S5). In the EAS and AFR populations, the results are similar to what we observed in figure S9: all the inbreeding coefficients are mostly unbiased but exhibit large variance (especially in the EAS population, panels E-L, tables S2-S5). Concerning the WORLD population, the most efficient estimates of ID are obtained with $F_{\text{AS}}$ (except for the $LMM_{GCTA^u}$ model, panels M-P, tables S2-S5). With both `GCTA` GRM matrices, all $F$ results in large variance around $b$ estimates (panels O and P, tables S4 and S5).

To conclude, $LMM_{\text{AS}}$ correctly estimates ID with the DOM scenario, in particular with $F_{\text{AS}}$ . However, we observe very large variance with smaller sample sizes: with some replicates overlapping with 0 in the WORLD population and many replicates in both the EAS and AFR populations. This is probably because the sample sizes are too small to estimate ID in these two populations.

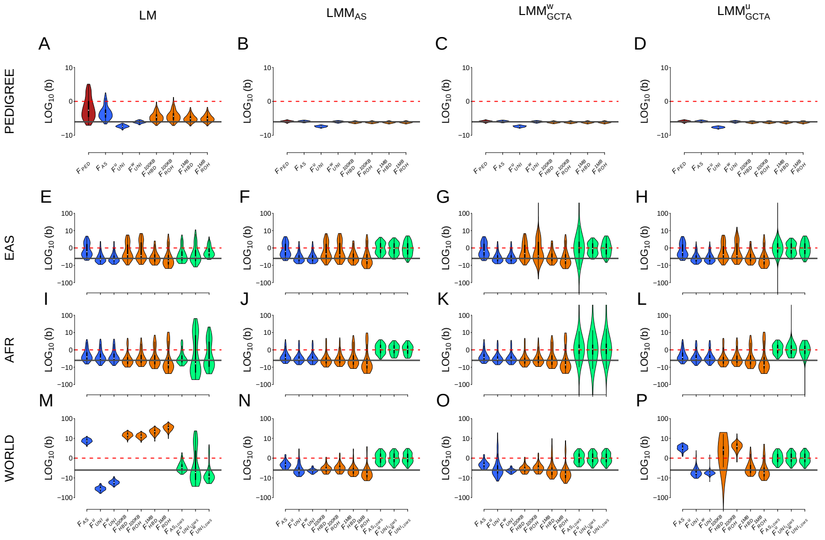


**Figure S11: Comparison of the estimation of inbreeding depression strength ($b$) among different $F$ estimates and models in four different populations with the DEMA scenario: effect sizes and dominance coefficients randomly assigned to causal markers and presence of DEMA.** Each column represents a regression model. The first column depicts the simple linear regression (panel A, E, I and M), the second column the linear mixed model with allele sharing GRM matrix as random factor (panel B, F, J and N), the third column the linear mixed model with the $GCTA^w$ relatedness matrix as random factor (panel C, G, K and O) and finally the forth column represents the linear mixed model with $GCTA^u$ relatedness matrix as random factor (panel D, H, L and P). The first row depicts the complete simulated population (11,924 individuals): PEDIGREE in panels A, B, C and D. Inbreeding estimates compared in these panels (A - D) are $F_{\text{PED}}$ , $F_{\text{AS}}$ , $F^u_{\text{UNI}}$ , $F^w_{\text{UNI}}$ , $F^{100KB}_{\text{HBD}}$ , $F^{100KB}_{\text{ROH}}$ , $F^{1MB}_{\text{HBD}}$ and $F^{1MB}_{\text{ROH}}$ .The last three rows are the populations from the 1,000 Genomes Project: EAS in panels E, F, G and H, AFR in panels I, J, K and L and WORLD in panels M, N, O and P. Inbreeding estimates compared in these panels (E - P) are $F_{\text{AS}}$ , $F^u_{\text{UNI}}$ , $F^w_{\text{UNI}}$ , $F^{100KB}_{\text{HBD}}$ , $F^{100KB}_{\text{ROH}}$ , $F^{1MB}_{\text{HBD}}$ , $F^{1MB}_{\text{ROH}}$ , $F_{\text{AS}_{\text{LDMS}}}$ , $F^u_{\text{UNI}_{\text{LDMS}}}$ and finally $F^w_{\text{UNI}_{\text{LDMS}}}$ . Violin plots represent the distribution of the inbreeding depression strength estimates (b) among the 100 replicates. The solid dark grey line is the true strength of ID ($b$ = -3). The dashed red line represents the absence of ID ($b$ = 0), meaning that we failed to detect ID in any replicate above this line. Note that all panels (A - P) are in $log_{10}$ scale. Also note that linear mixed models did not converge for some replicates (yielding estimated b values above 1000 or below -1000, not shown if outside the graph limits). **Percentages of replicates which did not converge:** panel G (EAS, $GCTA^w$ ): 1% for $F^{100KB}_{\text{ROH}}$ and 12% for $F_{\text{AS}_{\text{LDMS}}}$ ; panel H (EAS, $GCTA^u$ ): 2% for $F_{\text{AS}_{\text{LDMS}}}$ ; panel K (AFR, $GCTA^w$ ): 11% for $F_{\text{AS}_{\text{LDMS}}}$ , 17% for $F^u_{\text{UNI}_{\text{LDMS}}}$ and 14% for $F^w_{\text{UNI}_{\text{LDMS}}}$ ; panel L (AFR, $GCTA^u$ ): 1% for $F^u_{\text{UNI}_{\text{LDMS}}}$ and 1% for $F^w_{\text{UNI}_{\text{LDMS}}}$ ; panel P (WORLD, $GCTA^u$ ): 1% for $F^{100KB}_{\text{HBD}}$ .

Figure S11 shows the DEMA scenario (where both additive effects and dominance coefficients are independent of MAF but there is DEMA). Corresponding RMSE values can be found in tables S2-S5. With the simple LM and in the large PEDIGREE population, only $F_{\text{UNI}}$ and to a lesser extent IDB segments-based $F$ estimated with larger segments ($F_{\text{ROH}}^{1MB}$ and $F_{\text{HBD}}^{1MB}$ ) yield unbiased estimation of $b$ (panel A, table S2). Meanwhile, in the same large PEDIGREE population, the three mixed models allow efficient estimation of $b$ with all $F$ except $F_{\text{UNI}}^{u}$ , which overestimated the strength of ID (panels B-D, tables S3-S5). In the three populations from the 1,000 Genomes Project and especially in the EAS and WORLD populations, $F_{\text{AS}}$ is constantly overestimating the strength of ID (panels E-P, tables S2-S3). This is because DEMA is included in the simulations; $F_{\text{AS}}$ strongly correlates with the minor allele count (MAC) which makes it sensitive to DEMA [56]. Interestingly, LDMS stratification, allowed to get rid of the bias introduced by DEMA on $F_{\text{AS}}$ but it did not result in the most efficient estimation of $b$ (panel M, table S2-S5). Indeed, the lowest RMSE values are obtained with $F_{\text{UNI}}^{w}$ and both the $LMM_{\text{AS}}$ and $LMM_{GCTA^w}$ GRM emphasising that $F_{\text{UNI}}^{w}$ is robust to DEMA (tables S2-S5). With both $LMM_{\text{AS}}$ and $LMM_{GCTA^w}$ GRMs, IBD segments-based (especially based on larger segments: i.e. $F_{\text{ROH}}^{1MB}$ and $F_{\text{HBD}}^{1MB}$ ) yield results just behind $F_{\text{UNI}}^{w}$ (tables S2-S5).

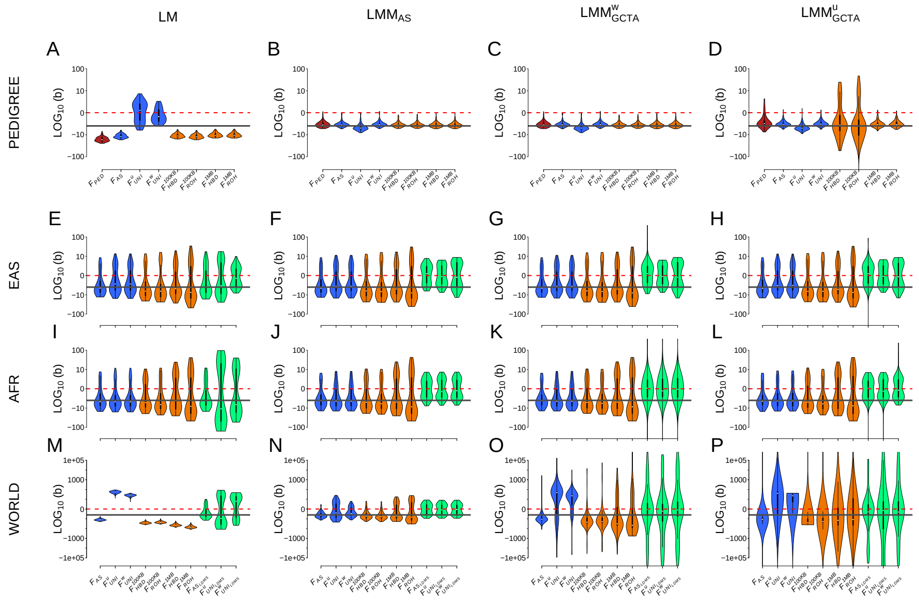


**Figure S12: Comparison of the estimation of inbreeding depression strength ($b$) among different $F$ estimates and models in four different populations with the ADD & DOM scenario: effect sizes and dominance coefficients assigned proportional to the MAF of causal markers and no DEMA.** Each column represents a regression model. The first column depicts the simple linear regression (panel A, E, I and M), the second column the linear mixed model with allele sharing GRM matrix as random factor (panel B, F, J and N), the third column the linear mixed model with the $GCTA^w$ relatedness matrix as random factor (panel C, G, K and O) and finally the forth column represents the linear mixed model with $GCTA^u$ relatedness matrix as random factor (panel D, H, L and P). The first row depicts the complete simulated population (11,924 individuals): PEDIGREE in panels A, B, C and D. Inbreeding estimates compared in these panels (A - D) are $F_{\text{PED}}$ , $F_{\text{AS}}$ , $F^u_{\text{UNI}}$ , $F^w_{\text{UNI}}$ , $F^{100KB}_{\text{HBD}}$ , $F^{100KB}_{\text{ROH}}$ , $F^{1MB}_{\text{HBD}}$ and $F^{1MB}_{\text{ROH}}$ .The last three rows are the populations from the 1,000 Genomes Project: EAS in panels E, F, G and H, AFR in panels I, J, K and L and WORLD in panels M, N, O and P. Inbreeding estimates compared in these panels (E - P) are $F_{\text{AS}}$ , $F^u_{\text{UNI}}$ , $F^w_{\text{UNI}}$ , $F^{100KB}_{\text{HBD}}$ , $F^{100KB}_{\text{ROH}}$ , $F^{1MB}_{\text{HBD}}$ , $F^{1MB}_{\text{ROH}}$ , $F_{\text{AS}_{\text{LDMS}}}$ , $F^u_{\text{UNI}_{\text{LDMS}}}$ and finally $F^w_{\text{UNI}_{\text{LDMS}}}$ . Violin plots represent the distribution of the inbreeding depression strength estimates (b) among the 100 replicates. The solid dark grey line is the true strength of ID ($b =$ -3). The dashed red line represents the absence of ID ($b = 0$), meaning that we failed to detect ID in any replicate above this line. Note that all panels (A - P) are in $log_{10}$ scale. Also note that linear mixed models did not converge for some replicates (yielding estimated b values above 1000 or below -1000, not shown if outside the graph limits). **Percentages of replicates which did not converge:** panel G (EAS, $GCTA^w$ ): 1% for $F_{\text{AS}_{\text{LDMS}}}$ ; panel H (EAS, $GCTA^u$ ): 1% for $F_{\text{AS}_{\text{LDMS}}}$ ; panel K (AFR, $GCTA^w$ ): 5% for $F_{\text{AS}_{\text{LDMS}}}$ , 5% for $F^u_{\text{UNI}_{\text{LDMS}}}$ and 7% for $F^w_{\text{UNI}_{\text{LDMS}}}$ ; panel L (AFR, $GCTA^u$ ): 1% for $F_{\text{AS}_{\text{LDMS}}}$ and 1% for $F^u_{\text{UNI}_{\text{LDMS}}}$ ; panel O (WORLD, $GCTA^w$ ): 1% for $F_{\text{AS}}$ , 5% for $F^u_{\text{UNI}}$ , 3% for $F^w_{\text{UNI}}$ , 5% for $F^{100KB}_{\text{HBD}}$ , 8% for $F^{100KB}_{\text{ROH}}$ , 6% for $F^{1MB}_{\text{HBD}}$ , 3% for $F^{1MB}_{\text{ROH}}$ , 28% for $F_{\text{AS}_{\text{LDMS}}}$ , 32% for $F^u_{\text{UNI}_{\text{LDMS}}}$ and 33% for $F^w_{\text{UNI}_{\text{LDMS}}}$ ; panel P (WORLD, $GCTA^u$ ): 4% for $F_{\text{AS}}$ , 9% for $F^u_{\text{UNI}}$ , 4% for $F^w_{\text{UNI}}$ , 3% for $F^{100KB}_{\text{HBD}}$ , 6% for $F^{100KB}_{\text{ROH}}$ , 11% for $F^{1MB}_{\text{HBD}}$ , 11% for $F^{1MB}_{\text{ROH}}$ , 22% for $F_{\text{AS}_{\text{LDMS}}}$ , 28% for $F^u_{\text{UNI}_{\text{LDMS}}}$ and 32% for $F^w_{\text{UNI}_{\text{LDMS}}}$ ; panel P (POLYPED, $GCTA^u$ ): 1% for $F^{100KB}_{\text{ROH}}$ .

Figure S12 shows the ADD & DOM scenario (where both additive effects and dominance coefficients are proportional to MAF and there is no DEMA). Corresponding RMSE values can be found in tables S2-S5. In the complete PEDIGREE population and with the simple LM, the strength of ID is underestimated by both $F_{\text{UNI}}$ and overestimated by all other $F$ (panel A, table S2). Similarly to what was observed with the previous scenarios, both $LMM_{\text{AS}}$ and $LMM_{GCTA^w}$ models allow efficient estimation of $b$ with all the $F$ (panels B and C, tables S3-S4). However, the $LMM_{GCTA^u}$ model results in biased estimation of $b$ with the short IBD segments-based $F$ ($F_{\text{ROH}}^{100KB}$ and $F_{\text{HBD}}^{100KB}$ , panel D, table S5). Similarly to what was observed before, we see no difference between the four models (except for the LDMS-based $F$) in both the EAS and AFR homogeneous populations (panels E-L, tables S2-S5). In the highly structured WORLD population however, the lowest RMSE values are obtained with the $LMM_{\text{AS}}$ model and especially with $F_{\text{AS}}$ (closely followed by $F_{\text{UNI}}^{w}$ , panel N, tables S2-S5). This is because DEMA is not included in this model and would strongly bias $b$ estimation with $F_{\text{AS}}$ . RMSE values are much larger for all $F$ with both `GCTA` matrices (panels O and P, tables S2-S5).

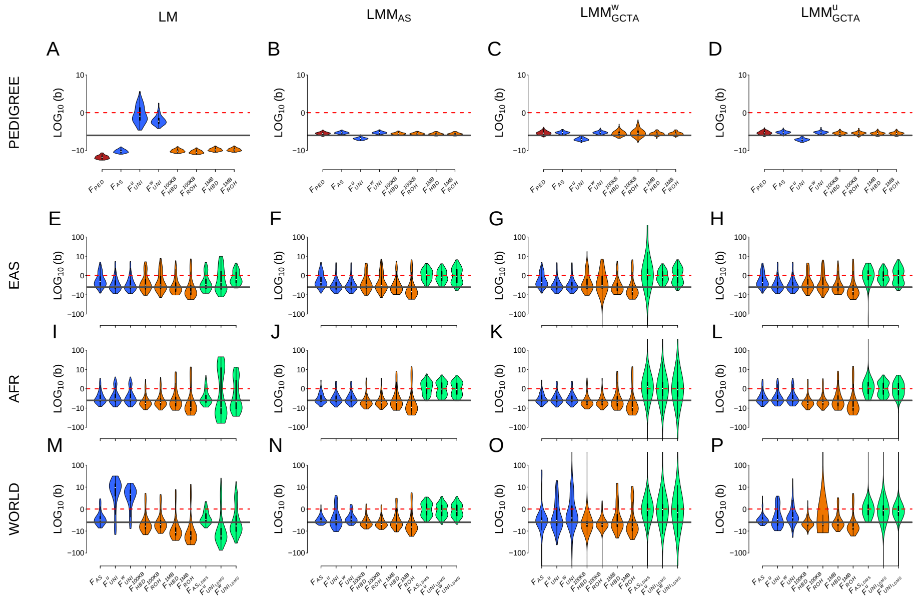


**Figure S13: Comparison of the estimation of inbreeding depression strength ($b$) among different $F$ estimates and models in four different populations with the ADD & DEMA scenario: effect sizes assigned proportional to the MAF of causal markers, dominance coefficients randomly assigned to causal markers and presence of DEMA.** Each column represents a regression model. The first column depicts the simple linear regression (panel A, E, I and M), the second column the linear mixed model with allele sharing GRM matrix as random factor (panel B, F, J and N), the third column the linear mixed model with the $GCTA^w$ relatedness matrix as random factor (panel C, G, K and O) and finally the forth column represents the linear mixed model with $GCTA^u$ relatedness matrix as random factor (panel D, H, L and P). The first row depicts the complete simulated population (11,924 individuals): PEDIGREE in panels A, B, C and D. Inbreeding estimates compared in these panels (A - D) are $F_{\text{PED}}$ , $F_{\text{AS}}$ , $F^u_{\text{UNI}}$ , $F^w_{\text{UNI}}$ , $F^{100KB}_{\text{HBD}}$ , $F^{100KB}_{\text{ROH}}$ , $F^{1MB}_{\text{HBD}}$ and $F^{1MB}_{\text{ROH}}$ .The last three rows are the populations from the 1,000 Genomes Project: EAS in panels E, F, G and H, AFR in panels I, J, K and L and WORLD in panels M, N, O and P. Inbreeding estimates compared in these panels (E - P) are $F_{\text{AS}}$ , $F^u_{\text{UNI}}$ , $F^w_{\text{UNI}}$ , $F^{100KB}_{\text{HBD}}$ , $F^{100KB}_{\text{ROH}}$ , $F^{1MB}_{\text{HBD}}$ , $F^{1MB}_{\text{ROH}}$ , $F_{\text{AS}_{\text{LDMS}}}$ , $F^u_{\text{UNI}_{\text{LDMS}}}$ and finally $F^w_{\text{UNI}_{\text{LDMS}}}$ . Violin plots represent the distribution of the inbreeding depression strength estimates (b) among the 100 replicates. The solid dark grey line is the true strength of ID ($b$ = -3). The dashed red line represents the absence of ID ($b = 0$), meaning that we failed to detect ID in any replicate above this line. Note that all panels (A - P) are in $log_{10}$ scale. Also note that linear mixed models did not converge for some replicates (yielding estimated b values above 1000 or below -1000, not shown if outside the graph limits). **Percentages of replicates which did not converge:** panel G (EAS, $GCTA^w$ ): 1% for $F^{100KB}_{\text{ROH}}$ and 16% for $F_{\text{AS}_{\text{LDMS}}}$ ; panel H (EAS, $GCTA^u$ ): 2% for $F_{\text{AS}_{\text{LDMS}}}$ ; panel K (AFR, $GCTA^w$ ): 13% for $F_{\text{AS}_{\text{LDMS}}}$ , 13% for $F^u_{\text{UNI}_{\text{LDMS}}}$ and 15% for $F^w_{\text{UNI}_{\text{LDMS}}}$ ; panel L (AFR, $GCTA^u$ ): 2% for $F_{\text{AS}_{\text{LDMS}}}$ and 1% for $F^w_{\text{UNI}_{\text{LDMS}}}$ ; panel O (WORLD, $GCTA^w$ ): 1% for $F_{\text{AS}}$ , 2% for $F^w_{\text{UNI}}$ , 3% for $F^{100KB}_{\text{HBD}}$ , 1% for $F^{100KB}_{\text{ROH}}$ , 7% for $F_{\text{AS}_{\text{LDMS}}}$ , 14% for $F^u_{\text{UNI}_{\text{LDMS}}}$ and 19% for $F^w_{\text{UNI}_{\text{LDMS}}}$ ; panel P (WORLD, $GCTA^u$ ): 1% for $F^{100KB}_{\text{ROH}}$ , 2% for $F_{\text{AS}_{\text{LDMS}}}$ , 4% for $F^u_{\text{UNI}_{\text{LDMS}}}$ and 2% for $F^w_{\text{UNI}_{\text{LDMS}}}$ .

Figure S13 shows the results of ID strength estimation for the ADD & DEMA scenario (when the additive effects are inversely proportional to MAF, the dominance effects are independent of MAF and there is presence of DEMA). Corresponding RMSE values can be found in tables S2-S5. Similarly to what was observed in the previous figures with the simple LM, the strength of ID is underestimated by both $F_{\text{UNI}}$ and overestimated by all other $F$ in the complete PEDIGREE population (panel A, table S2). The $LMM_{\text{AS}}$ and $LMM_{GCTA^u}$ models allow for a efficient estimation of $b$ with all $F$ except $F^u_{\text{UNI}}$ which results in a lightly overestimated estimation of $b$ (panels B and D, tables S2-S5). As for the $LMM_{GCTA^w}$ model, $F^u_{\text{UNI}}$ overestimates the strength of ID and the variance among $b$ estimates was larger with both short IBD segments-based $F$ ($F^{100KB}_{\text{ROH}}$ and $F^{100KB}_{\text{HBD}}$ , panel C, table S5). In both the homogeneous EAS and AFR populations there are not much differences among the models (panels E - L, tables S2-S5). However, in the WORLD population, the smallest RMSE are obtained with the $LMM_{\text{AS}}$ model (panel N, table S2-S5).

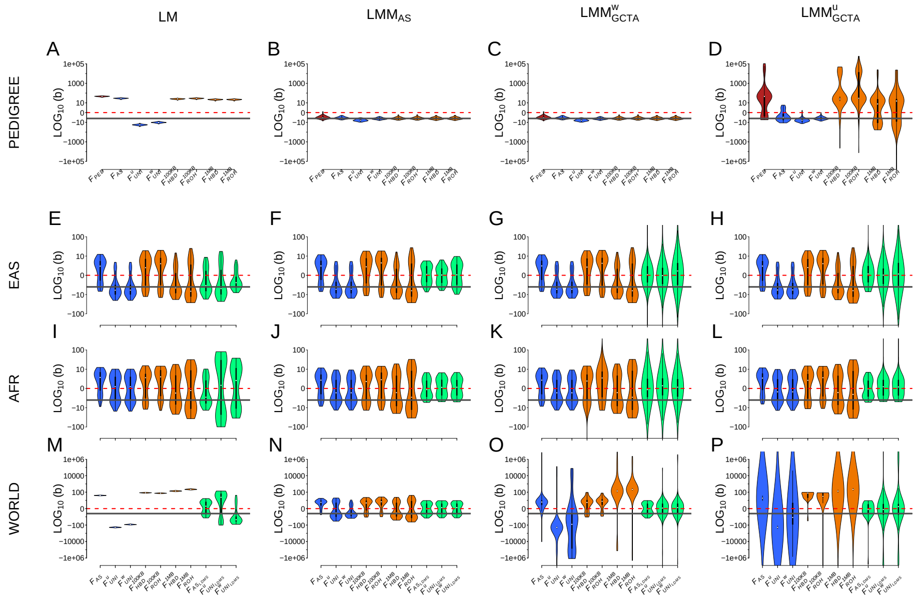


**Figure S14: Comparison of the estimation of inbreeding depression strength ($b$) among different $F$ estimates and models in four different populations with the DOM & DEMA scenario: effect sizes randomly assigned to causal markers, dominance coefficients assigned proportional to the MAF of causal markers and presence of DEMA.** Each column represents a regression model. The first column depicts the simple linear regression (panel A, E, I and M), the second column the linear mixed model with allele sharing GRM matrix as random factor (panel B, F, J and N), the third column the linear mixed model with the $GCTA^w$ relatedness matrix as random factor (panel C, G, K and O) and finally the forth column represents the linear mixed model with $GCTA^u$ relatedness matrix as random factor (panel D, H, L and P). The first row depicts the complete simulated population (11,924 individuals): PEDIGREE in panels A, B, C and D. Inbreeding estimates compared in these panels (A - D) are $F_{\text{PED}}$ , $F_{\text{AS}}$ , $F^u_{\text{UNI}}$ , $F^w_{\text{UNI}}$ , $F^{100KB}_{\text{HBD}}$ , $F^{100KB}_{\text{ROH}}$ , $F^{1MB}_{\text{HBD}}$ and $F^{1MB}_{\text{ROH}}$ .The last three rows are the populations from the 1,000 Genomes Project: EAS in panels E, F, G and H, AFR in panels I, J, K and L and WORLD in panels M, N, O and P. Inbreeding estimates compared in these panels (E - P) are $F_{\text{AS}}$ , $F^u_{\text{UNI}}$ , $F^w_{\text{UNI}}$ , $F^{100KB}_{\text{HBD}}$ , $F^{100KB}_{\text{ROH}}$ , $F^{1MB}_{\text{HBD}}$ , $F^{1MB}_{\text{ROH}}$ , $F_{\text{AS}_{\text{LDMS}}}$ , $F^u_{\text{UNI}_{\text{LDMS}}}$ and finally $F^w_{\text{UNI}_{\text{LDMS}}}$ . Violin plots represent the distribution of the inbreeding depression strength estimates (b) among the 100 replicates. The solid dark grey line is the true strength of ID ($b$ = -3). The dashed red line represents the absence of ID ($b = 0$), meaning that we failed to detect ID in any replicate above this line. Note that all panels (A - P) are in $log_{10}$ scale. Also note that linear mixed models did not converge for some replicates (yielding estimated b values above 1000 or below -1000, not shown if outside the graph limits). **Percentages of replicates which did not converge:** panel D (PEDIGREE, $GCTA^u$ ): 10% for $F_{\text{PED}}$ , 25% $F^{100KB}_{\text{HBD}}$ , 31% $F^{100KB}_{\text{ROH}}$ , 4% for $F^{1MB}_{\text{HBD}}$ and 9% for $F^{1MB}_{\text{ROH}}$ ; panel G (EAS, $GCTA^w$ ): 3% for $F_{\text{AS}_{\text{LDMS}}}$ , 8% for $F^u_{\text{UNI}_{\text{LDMS}}}$ and 8% for $F^w_{\text{UNI}_{\text{LDMS}}}$ ; panel H (EAS, $GCTA^u$ ): 3% for $F_{\text{AS}_{\text{LDMS}}}$ , 10% for $F^u_{\text{UNI}_{\text{LDMS}}}$ and 10% for $F^w_{\text{UNI}_{\text{LDMS}}}$ ; panel K (AFR, $GCTA^w$ ): 1% for $F^{100KB}_{\text{HBD}}$ , 2% for $F^{100KB}_{\text{ROH}}$ , 15% for $F_{\text{AS}_{\text{LDMS}}}$ , 9% for $F^u_{\text{UNI}_{\text{LDMS}}}$ and 12% for $F^w_{\text{UNI}_{\text{LDMS}}}$ ; panel L (AFR, $GCTA^u$ ): 1% for $F^u_{\text{UNI}_{\text{LDMS}}}$ and 1% for $F^w_{\text{UNI}_{\text{LDMS}}}$ ; panel O (WORLD, $GCTA^w$ ): 4% for $F_{\text{AS}}$ , 15% for $F^u_{\text{UNI}}$ , 45% for $F^w_{\text{UNI}}$ , 16% for $F^{1MB}_{\text{HBD}}$ , 12% for $F^{1MB}_{\text{ROH}}$ , 3% for $F^u_{\text{UNI}_{\text{LDMS}}}$ and 3% for $F^w_{\text{UNI}_{\text{LDMS}}}$ ; panel P (WORLD, $GCTA^u$ ): 15% for $F_{\text{AS}}$ , 28% for $F^u_{\text{UNI}}$ , 26% for $F^w_{\text{UNI}}$ , 1% for $F^{100KB}_{\text{ROH}}$ , 14% for $F^{1MB}_{\text{HBD}}$ , 10% for $F^{1MB}$[illegible] , 2% for $F_{\text{AS}}$[illegible] , 13% for $F^u$[illegible] and 16% for $F^w$[illegible]

Figure S14 shows the results of ID strength estimation for the DOM & DEMA scenario (when the additive effects are randomly assigned, the dominance effects are inversely proportional to MAF and there is presence of DEMA). Corresponding RMSE values can be found in tables S2-S5. Interestingly, we find the opposite of previous findings with LM in the entire PEDIGREE population: both $F_{\mathrm{UNI}}$ overestimate the strength of ID whereas all the other $F$ underestimate it (panel A). Both $LMM_{\mathrm{AS}}$ and $LMM_{GCTA^w}$ models allow for efficient estimation of $b$ with all the $F$ (panels B and C, tables S3-S4). With the $LMM_{GCTA^u}$ however, the estimation of $b$ for $F_{\mathrm{PED}}$ and all IBD segments-based $F$ was really poor. There is no difference between the four regression models for both homogeneous populations (panels E - L, tables S2-S5). Concerning the WORLD population, the smallest RMSE values were only obtained with the $LMM_{\mathrm{AS}}$ model, both GCTA matrices resulted in very large variances among $b$ estimates (panels M-P, tables S2-S5).

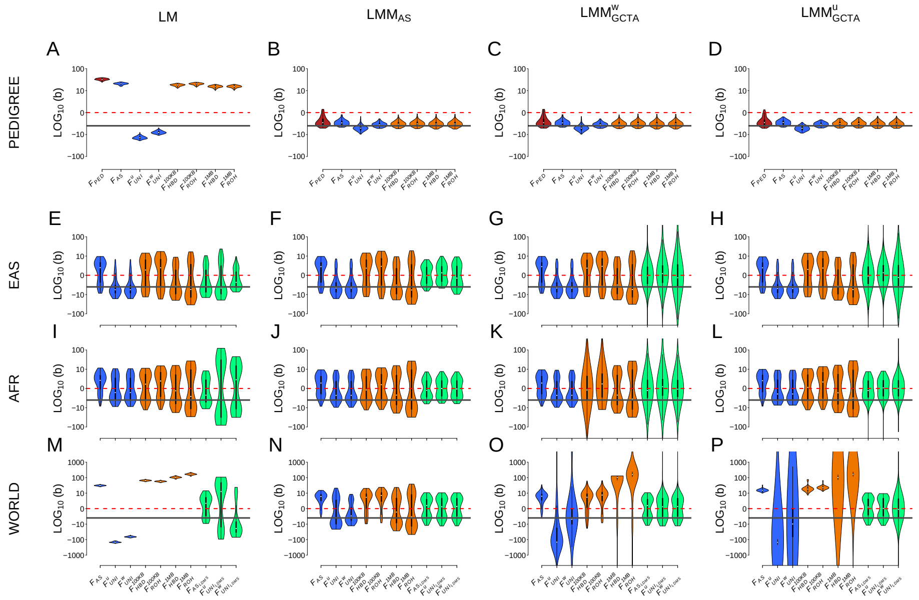


**Figure S15: Comparison of the estimation of inbreeding depression strength ($b$) among different $F$ estimates and models in four different populations with the ADD & DOM & DEMA scenario: effect sizes and dominance coefficients assigned proportional to the MAF of causal markers and presence of DEMA.** Each column represents a regression model. The first column depicts the simple linear regression (panel A, E, I and M), the second column the linear mixed model with allele sharing GRM matrix as random factor (panel B, F, J and N), the third column the linear mixed model with the $GCTA^w$ relatedness matrix as random factor (panel C, G, K and O) and finally the forth column represents the linear mixed model with $GCTA^u$ relatedness matrix as random factor (panel D, H, L and P). The first row depicts the complete simulated population (11,924 individuals): PEDIGREE in panels A, B, C and D. Inbreeding estimates compared in these panels (A - D) are $F_{\text{PED}}$ , $F_{\text{AS}}$ , $F^u_{\text{UNI}}$ , $F^w_{\text{UNI}}$ , $F^{100KB}_{\text{HBD}}$ , $F^{100KB}_{\text{ROH}}$ , $F^{1MB}_{\text{HBD}}$ and $F^{1MB}_{\text{ROH}}$ .The last three rows are the populations from the 1,000 Genomes Project: EAS in panels E, F, G and H, AFR in panels I, J, K and L and WORLD in panels M, N, O and P. Inbreeding estimates compared in these panels (E - P) are $F_{\text{AS}}$ , $F^u_{\text{UNI}}$ , $F^w_{\text{UNI}}$ , $F^{100KB}_{\text{HBD}}$ , $F^{100KB}_{\text{ROH}}$ , $F^{1MB}_{\text{HBD}}$ , $F^{1MB}_{\text{ROH}}$ , $F_{\text{AS}_{\text{LDMS}}}$ , $F^u_{\text{UNI}_{\text{LDMS}}}$ and finally $F^w_{\text{UNI}_{\text{LDMS}}}$ . Violin plots represent the distribution of the inbreeding depression strength estimates (b) among the 100 replicates. The solid dark grey line is the true strength of ID ($b$ = -3). The dashed red line represents the absence of ID ($b$ = 0), meaning that we failed to detect ID in any replicate above this line. Note that all panels (A - P) are in $log_{10}$ scale. Also note that linear mixed models did not converge for some replicates (yielding estimated b values above 1000 or below -1000, not shown if outside the graph limits). **Percentages of replicates which did not converge:** panel G (EAS, $GCTA^w$ ): 4% for $F_{\text{AS}_{\text{LDMS}}}$ , 7% for $F^u_{\text{UNI}_{\text{LDMS}}}$ and 13% for $F^w_{\text{UNI}_{\text{LDMS}}}$ ; panel H (EAS, $GCTA^u$ ): 6% for $F_{\text{AS}_{\text{LDMS}}}$ , 6% for $F^u_{\text{UNI}_{\text{LDMS}}}$ and 8% for $F^w_{\text{UNI}_{\text{LDMS}}}$ ; panel K (AFR, $GCTA^w$ ): 3% for $F^{100KB}_{\text{HBD}}$ , 1% for $F^{100KB}_{\text{ROH}}$ , 12% for $F_{\text{AS}_{\text{LDMS}}}$ , 15% for $F^u_{\text{UNI}_{\text{LDMS}}}$ and 12% for $F^w_{\text{UNI}_{\text{LDMS}}}$ ; panel L (AFR, $GCTA^u$ ): 1% for $F_{\text{AS}_{\text{LDMS}}}$ , 1% for $F^u_{\text{UNI}_{\text{LDMS}}}$ and 1% for $F^w_{\text{UNI}_{\text{LDMS}}}$ ; panel O (WORLD, $GCTA^w$ ): 9% for $F^u_{\text{UNI}}$ , 21% for $F^w_{\text{UNI}}$ , 3% for $F^{1MB}_{\text{HBD}}$ , 6% for $F^{1MB}_{\text{ROH}}$ , 1% for $F^u_{\text{UNI}_{\text{LDMS}}}$ and 1% for $F^w_{\text{UNI}_{\text{LDMS}}}$ ; panel P (WORLD, $GCTA^u$ ): 14% for $F^u_{\text{UNI}}$ , 24% for $F^w_{\text{UNI}}$ , 13% for $F^{1MB}_{\text{HBD}}$ , 6% for $F^{1MB}_{\text{ROH}}$ and 3% for $F^w_{\text{UNI}_{\text{LDMS}}}$ .

Figure S15 shows the results of ID strength estimation for the ADD & DOM & DEMA scenario (when both the additive effects and dominance coefficients are inversely proportional to MAF and there is presence of DEMA). Corresponding RMSE values can be found in tables S2-S5. We find the same results as figure S14 with the LM in the complete PEDIGREE population: both $F_{\mathrm{UNI}}$ are overestimating the strength of ID while all the other $F$ are largely underestimating it (panel A), suggesting that this pattern is due to the combination of the dominance coefficients being proportional to MAF and the presence of DEMA scenarios. In this scenario however, the three mixed models allow an efficient estimation of $b$ with all the $F$ (panels B - D, tables S2-S5). There is no difference between the four models for both homogeneous populations (panels E - L, tables S2-S5). Concerning the WORLD population, the smallest RMSE values were achieved only with the $LMM_{\mathrm{AS}}$ model and more specifically with $F^{w}_{\mathrm{UNI}}$ (panelsM-Pm, tables S2-S5). The variance among $b$ estimates however are still fairly large, suggesting that we lack power (i.e. the sample size is too small). Finally, both `GCTA` matrices resulted in biased $b$ estimates (panels O and P, tables S4 and S5).

To summarize, we show that ADD, DOM and DEMA largely increase the variance around $b$ estimates (particularly when more than one parameter was used). $F^{u}_{\mathrm{UNI}}$ and all IBD segments-based $F$ were especially sensitive to the additive effect sizes being proportional to MAF. All $F$ resulted in less accurate estimation of $b$ when the dominance coefficients were proportional to MAF. Perhaps, this is because our current model is not accounting for dominance, despite knowing there is dominance. This problem might be solved by including an additional random factor with the dominance coefficients. Finally, $F_{\mathrm{AS}}$ and to a lesser extent $F^{u}_{\mathrm{UNI}}$ were strongly influenced by DEMA. This might be strange as Yengo et al. [56] showed that $F^{u}_{\mathrm{UNI}}$ was robust to DEMA but we show in figure S16 that it might because they filtered on MAF $< 0.05$ in their analyses.

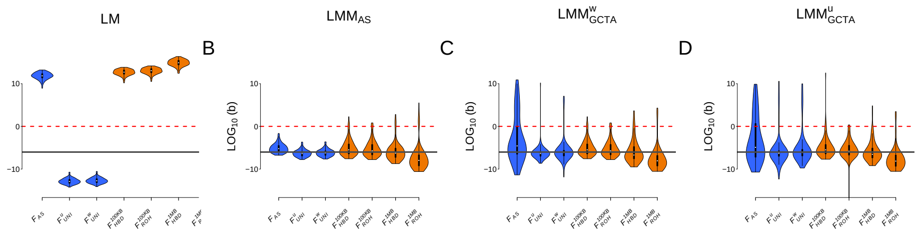


**Figure S16: Comparison of the estimation of inbreeding depression strength ($b$) among different $F$ estimates and models in the WORLD populations with the ADD & DOM & DEMA simulated scenario. SNPs have been filtered on $MAF > 0.05$:** panel A shows the simple linear model; panel B shows the linear mixed model with the allele sharing relatedness matrix as random factor; panel C shows the linear mixed model with $GCTA^w$ relatedness matrix as random factor; panel D shows the linear mixed model with $GCTA^u$ relatedness matrix as random factor. Violin plots represent the distribution of the inbreeding depression strength estimates (b) among the 100 replicates. The solid dark grey line is the true strength of ID ($b$ = -3). The dashed red line represents the absence of ID ($b$ = 0), meaning that we failed to detect ID in any replicate above this line.

Figure S16 shows the results of ID strength estimation for the ADD & DOM & DEMA scenario (when both the additive effects and dominance coefficients are inversely proportional to MAF and there is presence of DEMA) but where SNPs have been filtered on MAF, excluding all SNPs with MAF $<$ 0.05. We did this because the difference between $F^w_{\text{UNI}}$ and $F^u_{\text{UNI}}$ is the weight given to rare and common alleles. Consequently, we first filtered on MAF and then ran the same analyses ($F$ and GRMs estimation, as well as inbreeding depression simulations) on the WORLD population. We see that there is no difference between $F^w_{\text{UNI}}$ and $F^u_{\text{UNI}}$ when rare alleles are removed. This is because $F^u_{\text{UNI}}$ uses the average of ratios, which results in loci with small MAF strongly influencing the outcome. When these rare loci are filtered out, the estimated $F$ is no longer biased. This explains why Yengo *et al.* (2017) [56] found that $F^u_{\text{UNI}}$ was the best $F$ for quantifying inbreeding depression with an homogeneous subset of the UK bio bank dataset: they filtered on $MAF > 0.05$ leading to $F^u_{\text{UNI}}$ estimation not being influenced by rare alleles with strong additive and/or dominance effect sizes.

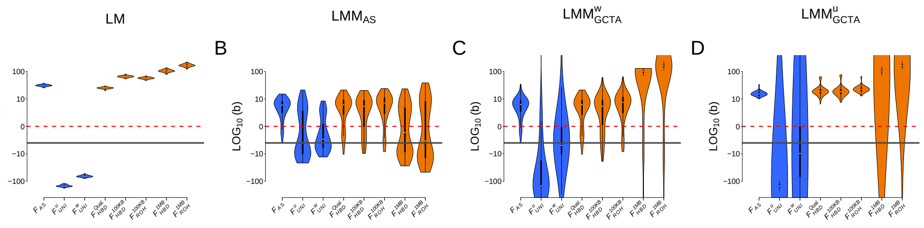


**Figure S17: Comparison of the estimation of inbreeding depression strength ($b$) among different $F$ estimates (including $F_{HBD.QUAL}$ with no minimum size threshold) and statistical models in the WORLD populations with the ADD & DOM & DEMA scenario: effect sizes and dominance coefficients assigned proportional to the MAF of causal markers and presence of DEMA.** Panel A shows the simple linear model; panel B shows the linear mixed model with the allele sharing relatedness matrix as random factor; panel C shows the linear mixed model with $GCTA^w$ relatedness matrix as random factor; panel D shows the linear mixed model with $GCTA^u$ relatedness matrix as random factor. Violin plots represent the distribution of the inbreeding depression strength estimates (b) among the 100 replicates. The solid dark grey line is the true strength of ID ($b$ = -3). The dashed red line represents the absence of ID ($b = 0$), meaning that we failed to detect ID in any replicate above this line.

Figure S17 shows the result of ID strength estimation for the ADD & DOM & DEMA scenario (when both the additive effects and dominance coefficients are inversely proportional to MAF and there is presence of DEMA) but with a new $F$: $F_{HBD.QUAL}$ where IBD segments were infered using `BCFTools` and no size selection was performed. Indeed, since `BCFTools` is a model-based HBD calling approach, there is no mandatory length restriction. The results are similar to those obtained with $F_{\text{HBD}}^{100KB}$ .

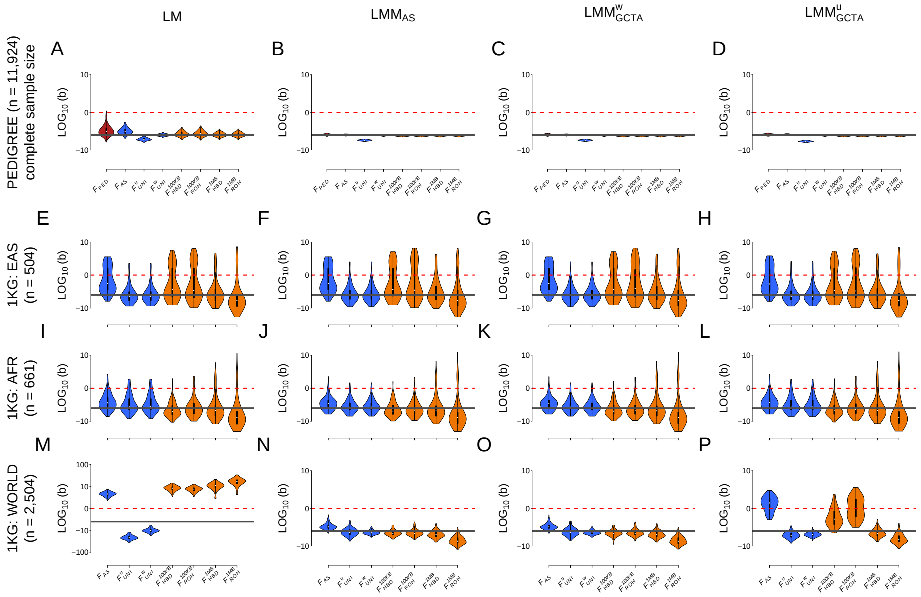


**Figure S18: Comparison of the estimation of inbreeding depression strength ($b$) among different $F$ estimates and models in four different populations with the ADD & DOM & DEMA scenario: effect sizes and dominance coefficients assigned proportional to the MAF of causal markers and presence of DEMA. Causal loci have been selected with intermediate frequencies:** $MAF > 0.1$ Each column represents a regression model. The first column depicts the simple linear regression (panel A, E, I and M), the second column the linear mixed model with allele sharing GRM matrix as random factor (panel B, F, J and N), the third column the linear mixed model with the $GCTA^w$ relatedness matrix as random factor (panel C, G, K and O) and finally the forth column represents the linear mixed model with $GCTA^u$ relatedness matrix as random factor (panel D, H, L and P). The first row depicts the complete simulated population (11,924 individuals): PEDIGREE in panels A, B, C and D. Inbreeding estimates compared in these panels (A - D) are $F_{\mathrm{PED}}$ , $F_{\mathrm{AS}}$ , $F^u_{\mathrm{UNI}}$ , $F^w_{\mathrm{UNI}}$ , $F^{100KB}_{\mathrm{HBD}}$ , $F^{100KB}_{\mathrm{ROH}}$ , $F^{1MB}_{\mathrm{HBD}}$ and $F^{1MB}_{\mathrm{ROH}}$ .The last three rows are the populations from the 1,000 Genomes Project: EAS in panels E, F, G and H, AFR in panels I, J, K and L and WORLD in panels M, N, O and P. Inbreeding estimates compared in these panels (E - P) are $F_{\mathrm{AS}}$ , $F^u_{\mathrm{UNI}}$ , $F^w_{\mathrm{UNI}}$ , $F^{100KB}_{\mathrm{HBD}}$ , $F^{100KB}_{\mathrm{ROH}}$ , $F^{1MB}_{\mathrm{HBD}}$ , $F^{1MB}_{\mathrm{ROH}}$ , $F_{\mathrm{AS_{LDMS}}}$ , $F^u_{\mathrm{UNI_{LDMS}}}$ and finally $F^w_{\mathrm{UNI_{LDMS}}}$ . Violin plots represent the distribution of the inbreeding depression strength estimates (b) among the 100 replicates. The solid dark grey line is the true strength of ID ($b$ = -3). The dashed red line represents the absence of ID $b = 0$), meaning that we failed to detect ID in any replicate above this line. Note that all panels (A - P) are in $log_{10}$ scale. Also note that all linear mixed models converged for all replicates.

As mentioned in the introduction, Alemu *et al.* (2021) [1] and Caballero *et al.* (2020) [7] showed that the best $F$ actually depends on the history of the population. Indeed, they showed that $F_{\text{ROH}}$ and $F_{\text{HBD}}$ and to a lesser extent $F_{\text{HOM}}$ were more efficient at quantifying homozygosity at loci with common alleles. On the contrary, $F^{u}_{\text{UNI}}$ was better at quantifying homozygosity at rare alleles. The authors propose that in populations with low effective sizes, the strength of selection is diminished which can lead to deleterious alleles reaching intermediate frequencies because of drift. This means that both $F_{\text{ROH}}$ and $F_{\text{HBD}}$ will perform better in such populations. On the contrary, in populations with large effective size, selection maintains deleterious alleles a low frequencies which explains why Yengo *et al.* (2017) found that $F_{\text{UNI}}$ was the best $F$ with the large UK biobank dataset. Consequently, we simulated an additional scenario where intermediate frequencies causal loci were selected on MAF $> 0.1$ (figure S18). Firstly, this greatly reduced the variance around $b$ estimates for all models and $F$. This suggests that it is much easier to detect inbreeding depression when the causal loci have higher frequencies. Secondly and as expected, it improved all IBD-segments based $F$ (both $F_{\text{HBD}}$ and both $F_{\text{ROH}}$ ) in all populations except $F^{100KB}_{\text{HBD}}$ and $F^{100KB}_{\text{ROH}}$ in the EAS population. This is because the EAS population has a small effective population size, resulting in large numbers of small IBD segments. Since inbreeding depression is mostly caused by recent coalescence events, including these smaller segments added noise in the models, leading to biased $b$ estimates. This is consistent with previous studies who showed that only including larger fragments resulted in better inbreeding depression estimates [7, 6]. Finally and surprisingly, excluding rare causal loci did not worsen $F^{w}_{\text{UNI}}$ estimation of $b$ as we would have expected. This might be because we did not explicitly simulate small populations, but simply selected common causal loci.

[16] Øystein Flagstad, Camilla Wikenros, Håkan Sand, Hans Ellegren, Marty Kardos, Mikael Åkesson, Olof Liberg, Pall Olason, Petter Wabakken, and Toby Fountain. Genomic consequences of intensive inbreeding in an isolated wolf population. *Nature Ecology & Evolution*, page 1, 2017.

[24] Marty Kardos, Yaolei Zhang, Kim M. Parsons, Yunga A, Hui Kang, Xun Xu, Xin Liu, Craig O. Matkin, Peijun Zhang, Eric J. Ward, M. Bradley Hanson, Candice Emmons, Michael J. Ford, Guangyi Fan, and Songhai Li. Inbreeding depression explains killer whale population dynamics. *Nature Ecology & Evolution*, 2023.

[33] Gustave Malécot. *Les mathématiques de l'hérédité*. Barnéoud frères, 1948.

[34] K. Martikainen, A. Sironen, and P. Uimari. Estimation of intrachromosomal inbreeding depression on female fertility using runs of homozygosity in Finnish Ayrshire cattle. *JOURNAL OF DAIRY SCIENCE*, 101(12):11097–11107, 2018.

[35] K. Martikainen, A. M. Tyriseva, K. Matilainen, J. Poso, and P. Uimari. Estimation of inbreeding depression on female fertility in the Finnish Ayrshire population. *JOURNAL OF ANIMAL BREEDING AND GENETICS*, 134(5):383–392, 2017.

[36] S. Mastrangelo, M. Tolone, R. Di Gerlando, L. Fontanesi, M. T. Sardina, and B. Portolano. Genomic inbreeding estimation in small populations: Evaluation of runs of homozygosity in three local dairy cattle breeds. *ANIMAL*, 10(5):746–754, 2016.

[37] Ruth McQuillan, Anne-Louise Leutenegger, Rehab Abdel-Rahman, Christopher S. Franklin, Marijana Pericic, Lovorka Barac-Lauc, Nina Smolej-Narancic, Branka Janicijevic, Ozren Polasek, Albert Tenesa, Andrew K. MacLeod, Susan M. Farrington, Pavao Rudan, Caroline Hayward, Veronique Vitart, Igor Rudan, Sarah H. Wild, Malcolm G. Dunlop, Alan F.

Wright, Harry Campbell, and James F. Wilson. Runs of homozygosity in European populations. *AMERICAN JOURNAL OF HUMAN GENETICS*, 83(3):359–372, 2008.

[38] R. Meyermans, W. Gorssen, N. Buys, and S. Janssens. How to study runs of homozygosity using PLINK? A guide for analyzing medium density SNP data in livestock and pet species. *BMC Genomics*, 21(1):94, 2020.

[39] Vagheesh Narasimhan, Petr Danecek, Aylwyn Scally, Yali Xue, Chris Tyler-Smith, and Richard Durbin. BCFtools/RoH: A hidden Markov model approach for detecting autozygosity from next-generation sequencing data. *Bioinformatics*, 32(11):1749–1751, 2016.

[40] Pirmin Nietlisbach, Stefanie Muff, Jane M. Reid, Michael C. Whitlock, and Lukas F. Keller. Nonequivalent lethal equivalents: Models and inbreeding metrics for unbiased estimation of inbreeding load. *Evolutionary Applications*, 0(0), 2019.

[41] Pascal A. Oltenacu and Bo Algers. Selection for Increased Production and the Welfare of Dairy Cows: Are New Breeding Goals Needed? *AMBIO: A Journal of the Human Environment*, 34(4):311–315, June 2005. Publisher: Royal Swedish Academy of Sciences.

[42] Elisa Peripolli, Nedenia Bonvino Stafuzza, Danisio Prado Munari, Andre Luis Ferreira Lima, Renato Irgang, Marco Antonio Machado, Joao Claudio do Carmo Panetto, Ricardo Vieira Ventura, Fernando Baldi, and Marcos Vinicius Gualberto Barbosa da Silva. Assessment of runs of homozygosity islands and estimates of genomic inbreeding in Gyr (Bos indicus) dairy cattle. *BMC GENOMICS*, 19, 2018.

[43] E. Postma, L. Martini, and P. Martini. Inbred women in a small and isolated Swiss village have fewer children. *Journal of Evolutionary Biology*, 23(7):1468–1474, 2010. _eprint: https://onlinelibrary.wiley.com/doi/pdf/10.1111/j.1420-9101.2010.02013.x.

[44] Jonathan K. Pritchard. Are Rare Variants Responsible for Susceptibility to Complex Diseases? *The American Journal of Human Genetics*, 69(1):124–137, July 2001.

[45] Jennie E. Pryce, Mekonnen Haile-Mariam, Michael E. Goddard, and Ben J. Hayes. Identification of genomic regions associated with inbreeding depression in Holstein and Jersey dairy cattle. *GENETICS SELECTION EVOLUTION*, 46, 2014.
